# Testing models of mRNA localization reveals robustness regulated by reducing transport between cells

**DOI:** 10.1101/533133

**Authors:** J. U. Harrison, R. M. Parton, I. Davis, R. E. Baker

## Abstract

Robust control of gene expression in both space and time is of central importance in the regulation of cellular processes, and for multicellular development. However, the mechanisms by which robustness is achieved are generally not identified or well understood. For example, mRNA localization by molecular-motor-driven transport is crucial for cell polarization in numerous contexts, but the regulatory mechanisms that enable this process to take place in the face of noise or significant perturbations are not fully understood. Here we use a combined experimental-theoretical approach to characterize the robustness of *gurken/TGF-alpha* mRNA localization in *Drosophila* egg chambers, where the oocyte and 15 surrounding nurse cells are connected in a stereotypic network via intracellular bridges known as ring canals. We construct a mathematical model that encodes simplified descriptions of the range of steps involved in mRNA localization, including production and transport between and within cells until the final destination in the oocyte. Using Bayesian inference, we calibrate this model using quantitative single molecule fluorescence in situ hybridization data. By analyzing both the steady state and dynamic behaviours of the model, we provide estimates for the rates of different steps of the localization process, as well as the extent of directional bias in transport through the ring canals. The model predicts that mRNA synthesis and transport must be tightly balanced to maintain robustness, a prediction which we tested experimentally using an over-expression mutant. Surprisingly, the over-expression mutant fails to display the anticipated degree of overaccumulation of mRNA in the oocyte predicted by the model. Through careful model-based analysis of quantitative data from the over-expression mutant we show evidence of saturation of transport of mRNA through ring canals. We conclude that this saturation engenders robustness of the localization process, in the face of significant variation in the levels of mRNA synthesis.

**Statement of significance:** For development to function correctly and reliably across a population, gene expression must be controlled robustly in a repeatable manner. How this robustness is achieved is not well understood. We use modelling to better study the localization of polarity determining transcripts (RNA) in fruit fly development. By calibrating our model with quantitative imaging data we are able to make experimentally testable predictions. Comparison of these predictions with data from a genetic mutant reveals evidence that saturation of RNA transport contributes to the robustness of RNA localization.

## 1 Introduction

Biological systems are constantly subjected to noise from both internal and external sources, and also to random mutations. It is crucial that key driving mechanisms are able to buffer against such insults, thereby enabling organisms to display phenotypes that are robust to perturbation. Localization of mRNA is a fundamental, and crucially important, mechanism that enables the polarization of cells such as oocytes and early embryos [1–3]. For example, the axes of *Drosophila* are established through the regulation of gradients in *bicoid, gurken, oskar* and *nanos* mRNA [4]. In fact, *Drosophila* and other species rely heavily on the asymmetric localization of mRNAs to coordinate early development processes both spatially and temporally. Further, mRNA localization has been observed in a variety of species and cell types, including *Drosophila* and *Xenopus* oocytes, neurons, chicken fibroblasts, yeast and bacteria [5–9], demonstrating that the process is ubiquitous and not limited to large cells. However, despite the widespread nature of mRNA localization, there is still much that we do not understand, in particular what ensures robustness. To this end, here we combine mathematical modelling with quantitative experimental data to investigate the regulatory mechanisms controlling mRNA localization, capturing, in particular, the roles of transport and production.

We concentrate on early development of the model organism *Drosophila*, where maternal mRNA prepatterns the oocyte [1–3]. Maternal mRNA is produced in the 15 nurse cells neighbouring the oocyte [10]; it is then packaged into complexes with various proteins [11] and transported through intracellular bridges, known as ring canals, into the oocyte [12]. Ring canals are formed by incomplete cell divisions [13], and result in the 15 nurse cells and the oocyte being connected in a characteristic pattern [14]. A schematic of the early *Drosophila* egg chamber is shown in Figure 1, together with a microscopy image showing the ring canals (highlighted with an actin (phalloidin Alexa488) marker), and a diagram illustrating the characteristic connections between nurse cells.

**Figure 1:**
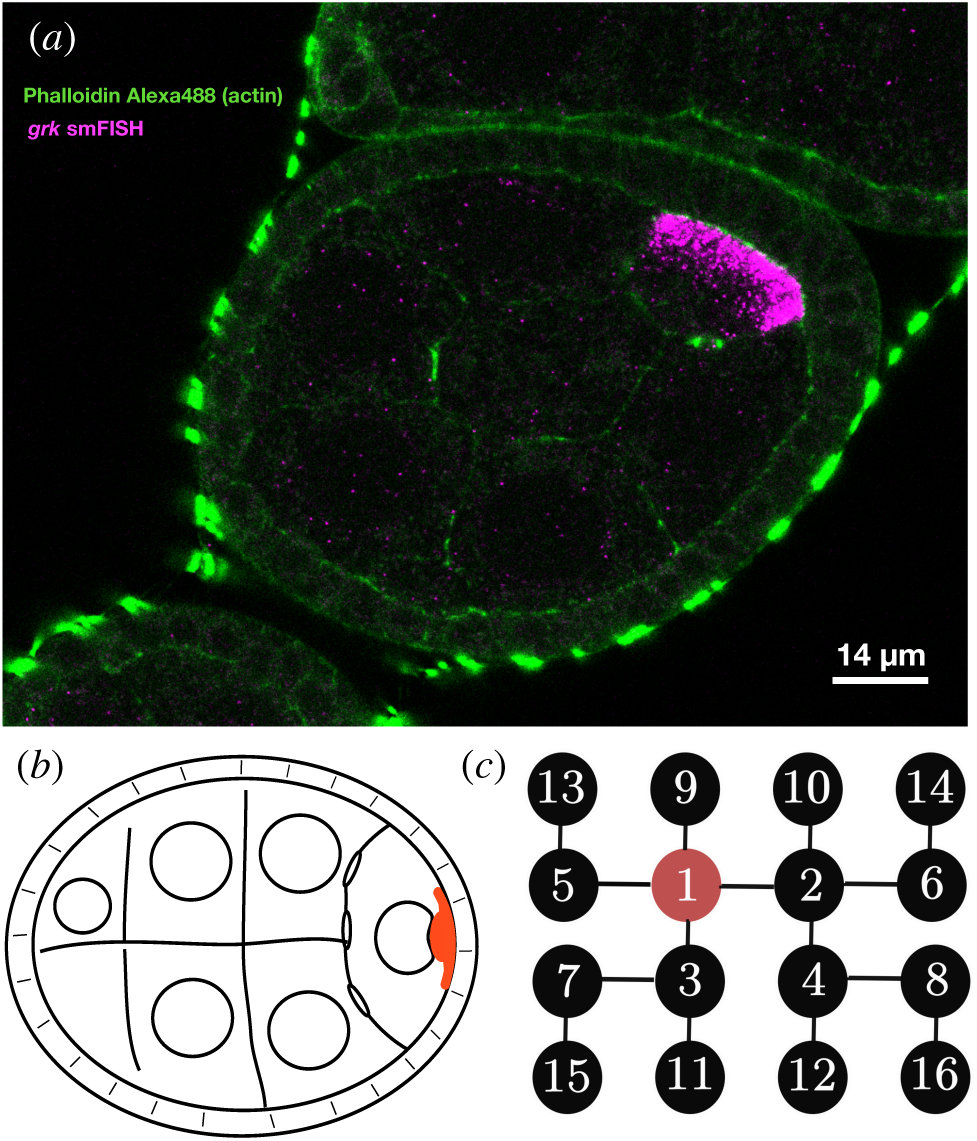
Stage 6 *Drosophila* egg chamber. (a) A microscopy image of a fixed egg chamber labelled in green with pahlloidin-alexa488 (actin) and in magenta with *grk* smFISH probes (stellaris, Cal Fluor 590). Localization of *gurken* RNA is observed at the posterior of the oocyte. Ring canals providing connections between nurse cells and to the oocyte can be seen highlighted by the actin marker. (b) Schematic of the early egg chamber. (c) Diagram of the connections between nurse cells and the ooctye via ring canals. The oocyte (red) is labeled 1 and the nurse cells are labeled 2 to 16.

In this work, we develop a coarse-grained ordinary differential equation (ODE) model of *gurken* (*gurken*) transport through the *Drosophila* egg chamber, including all 15 nurse cells and the developing oocyte, with terms representing both the production and transport of mRNA ^1^. Using Bayesian inference methods, we fit this simple model to quantitative imaging data obtained using smFISH (single molecule fluorescence *in situ* hybridization) [15, 16]. Combining quantitative modelling approaches with experimental data in this way allows us to explore underlying biological mechanisms through the generation of testable model predictions with appropriate quantification of uncertainties. In particular, it enables estimation of the mRNA production and transport rates in the model, and predicts that there is a tight regulatory balance between production and transport for the localization of *gurken* in the *Drosophila* egg chamber.

In the process of fitting the model to microscopy imaging data, we examine the formation of higher-order RNA-protein complexes in the oocyte. In the nurse cells, mRNA is assembled into complexes containing both mRNA and various proteins, and these RNA complexes are then transported into the ooctye. An important aspect of the transport process is that the RNA complexes are remodelled (so that more mRNA transcripts are contained in each complex) upon transit through the ring canals that connect the nurse cells to the oocyte [12, 17], leading to larger complexes in the oocyte [18]. We use quantitative data analysis to provide an estimate of the extent of this assembly of higher-order complexes for *gurken*.

In addition, we explore the question of whether transport of RNA complexes occurs unidirectionally or bidirectionally through ring canals in the *Drosophila* egg chamber. Some evidence that transport through ring canals is unidirectional has previously been provided [17]. However, since ring canals are small relative to the nurse cells, the passage of complexes through them is relatively difficult to observe *in vivo*. Our study offers further evidence in support of the hypothesis that transport through ring canals is strongly biased towards the oocyte, and provides quantification of this process within a model-based framework.

Finally, we use the coarse-grained ODE model to make testable predictions about the behaviour of an over-expression mutant with increased production of mRNA. We demonstrate strong agreement between the model predictions and observed data for nurse cells close to the oocyte, but find discrepancies for nurse cells far from the oocyte. Surprisingly, we find the numbers of complexes localized in the oocyte of the over-expression mutant are very similar to wild type, whereas the model predicts numbers should increase significantly. To probe the reasons for this disparity, we consider a suite of extended models incorporating inhomogeneous production, density dependent transport and crowding-induced blocking of transport through ring canals. We show, via statistical techniques that allow quantitative model comparison, that the crowding-induced blocking mechanism is best supported by the data, and that a model incorporating blocking is capable of producing distributions of RNA complexes similar to those observed experimentally. In strong support of this model-based prediction, we observe accumulation of complexes at ring canals in smFISH microscopy data for the over-expression mutant, consistent with this crowding-induced blocking mechanism.

Despite their increasing use across the biological sciences, mathematical and computational modelling approaches have not to-date been widely adopted in the study of mRNA localization. Trong et al. [19] have demonstrated how localization of mRNA can be achieved in a *Drosophila* oocyte using a partial differential equation (PDE) description of mRNA dynamics and a stochastic model of cytoskeleton dynamics. Ciocanel et al. [20] used a PDE model of RNA localization in *Xenopus* oocytes to quantify aspects of the transport process using FRAP (fluoresence recovery after photobleaching) data, and others have considered models of RNA transport to examine the behaviour of individual molecular motors [21, 22], or the cytoskeleton structure [23]. However, these works have focused on dynamics within a single cell, the oocyte, as has other work in model systems such as mating budding yeast and the neuronal growth cone [24, 25]. Recent work by Alsous et al. [26] has quantified collective growth and size control dynamics in *Drosophila* nurse cells. Similarly, our work considers dynamic behaviours in multiple cells. However, our focus is to examine the regulation and robustness of mRNA localization by considering a coarse-grained ODE model of the entire *Drosophila* egg chamber, and using it to interrogate quantitative experimental data. Further, our work highlights that by combining an incredibly simple mathematical model with quantitative experimental data one can draw relevant conclusions and make testable predictions about key biological processes that would not, otherwise, have been possible.

## 2 Methods and materials

### 2.1 Fly strains and tissue preparation

Stocks were raised on standard cornmeal-agar medium at 25°C. The wild type was Oregon R (OrR). Over expression of *gurken* RNA was obtained by crossing *UASp grk3A* (based upon genomic sequence DS02110, which includes the full 3’ and 5’ UTRs [27]) to maternal tubulin driver TubulinGal4-VP16. Tagging of *gurken* RNA for live imaging was achieved using the MS2-MCP(FP) system: *gurken*-(MS2)12 [28], P[*nos-NLS-MCP-mCherry*] [29]; P[*w+nanos-5’-NLS-MCP-Dendra2 K10-3’*] (Rippei Hayashi). GFP protein trap lines: Moesin::GFP [30]; Tau-GFP 65/167 (D. St Johnston, University of Cambridge). Oocytes were isolated for live imaging as described in Parton et al. [31], and for fixation and single molecule fluorescence in situ hybridization (smFISH), as described in Davidson et al. [32].

### 2.2 Single molecule FISH

smFISH detection of *gurken* RNA was performed as described in Davidson et al. [32]. Stellaris Olignoucleotide probes 20nt in length complementary to the *gurken* transcript (CG17610; 48 probes) conjugated to CAL Fluor Red 590 were obtained from Biosearch technologies. Labelled oocytes were mounted on slides in 70% Vectashield (Vector laboratories).

### 2.3 Fixed-cell imaging

Confocal imaging of fixed *Drosophila* oocytes was performed using an inverted Olympus FV3000 six laser line spectral confocal system fitted with high sensitivity gallium arsenide phosphide (GaAsP detectors) and using a 60x SI 1.3 NA lens. For smFISH detection of *gurken* transcripts, settings were optimized with a pinhole of 1.2 airy units and increasing laser power over the depth of the sample (between 30-50*µ*m from the cover slip) to compensate for signal attenuation.

### 2.4 Image analysis

Basic image handling and processing was carried out in FIJI (ImageJ V1.51d; *fiji.sc*, [33]). Image data was archived in OMERO (V5.3.5) [34]; image conversions were carried out using the BioFormats plugin in FIJI [35]. RNA particles in the nurse cells and oocyte are identified and located using the FISH QUANT software [36]. We combine use of this software with manual segmentation of the nurse cells and oocyte performed in three dimensions using a GUI (graphical user interface) designed for the annotation of training and evaluation data for machine learning via software available from QBrain [37]. This enables quantification of the total numbers of complexes observed in each cell. Identification of cells within the egg chamber is performed by counting the number of ring canals connecting each cell, and which cells these connect. Data is processed from *n* = 16 egg chambers between stage 5 and stage 8 of oogenesis. Since the RNA complexes are sparse in the nurse cells, using counts of particles is a valid method of quantifying the total RNA expression [38].

### 2.5 Identification of developmental time via an exponential growth model

To identify a precise time variable for each egg chamber, we apply an exponential growth model to measurements of the area of the median section. Effectively this allows us to use log(*A*), where *A* is the area of the median section, as a rescaled variable for the timescale of development, by fitting the following model by linear regression: log(*A*) = log(*A*_0_) + *τt*, where the intercept *t*_0_ = log(*A*_0_) corresponds to the start of *gurken* mRNA production, and *τ* gives a rescaling of time. To fit this model, we use data from Shimada et al. [39] relating the developmental stage to the area of egg chambers and take the time point corresponding to the midpoint of each stage as an estimate of its age (Figure S1). This approach is supported by the results of Jia et al. [40], who use log *A* in a similar way to assist with automatic stage identification for *Drosophila* egg chambers.

### 2.6 Implementation of Monte Carlo methods

We use the open source software package Stan *mc - stan.org* (version 2.17.0) [41] to perform MCMC (Markov chain Monte Carlo) sampling from the posterior distribution. Stan uses Hamiltonian Monte Carlo techniques to navigate the geometry of the posterior distribution efficiently [42], along with automatic differentiation to compute gradients. We run four chains in parallel for 2000 iterations with a burn in of 1000 iterations. Traceplots illustrating convergence of each of the chains are shown in Figure S2. Code to reproduce the analysis described in this work is available at https://github.com/shug3502/rstan_analysis.

### 2.7 Model comparison

We perform model comparison via leave-one-out cross validation [43–45]. From the family of models under consideration, we make testable predictions for the behaviour of a genetic over-expression mutant and compare these with experimental data. We compute pseudo-BMA+ (Bayesian Model Averaging) weights [46] for each model, where the largest weight will give the model closest in Kullback-Leibler divergence to the data generating model. However, these weights can potentially be misleading in the case where the true model is not contained in the set of models considered, as is the case here. In this ℳ-open case, stacking weights [46] offer a more conservative choice of models. We present pseudo-BMA+ and stacking weights, since both are informative. For further details see Supplementary Material Section K.

## 3. Results

### 3.1 A coarse-grained model for gurken mRNA localization

To investigate the mechanisms governing robustness of mRNA localization, we developed a minimal, compartment-based ODE model of *gurken* localization in a *Drosophila* egg chamber. We view each cell as a separate compartment and assume that RNA complexes are produced in each of the nurse cells [10]. We assume that there is transport of RNA complexes between cells that are connected by ring canals (Figure 1), and that there is negligible degradation (there is evidence that the majority of maternal mRNAs are stable during oogenesis over the timescales of interest [47, 48]).

The model can be written as

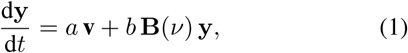

where the *i* ^th^ entry of the vector **y** is the number of RNA complexes in cell *i* (indexed as in Figure 1). RNA complexes are produced at constant rate *a >* 0 in units of [particles hr^−1^], and the vector **v** ∈ℝ^16^ lists the cells that produce mRNA; for the *Drosophila* egg chamber, where mRNA is produced in all cells except the ooctye, **v** = (0, 1, *…,* 1)^T^ since the oocyte is transcriptionally silent for most of oogenesis [49]. Transport of complexes between cells takes place at constant rate *b >* 0 in units of [hr^−1^]. The matrix **B** describes the network of connected nurse cells (so that entry (*i, j*) is non-zero only when cells *i* and *j* are connected by a ring canal). Transport bias (towards the oocyte) is represented by parameter *ν* so that complexes in a given nurse cell move towards the oocyte at relative rate 1 − *ν* and away from it at relative rate *ν* (Figure 4).The precise form of **B** is provided in Supplementary Material Section A. Note that a similar approach has been employed by Alsous et al. [26] to show that differences in cell sizes in the *Drosophila* egg chamber result from the characteristic cell network generated through incomplete divisions, and the resulting ring canal connections. As an initial condition, we assume there are no RNA complexes in any of the cells at time *t*_0_ (where *t*_0_ is determined from the linear model described in Section 2.5). The system of ODEs is solved numerically using the fourth order Runge-Kutta scheme implementation contained in the Boost C++ library [50].

Typical behaviour of the model is shown in Figure 2. Much of the biologically relevant behaviour occurs in the quasi-steady-state regime where we have linear growth in the number of RNA complexes in the oocyte, and a characteristic distribution of RNA complexes across the nurse cells. Typical dynamic behaviour for each nurse cell is shown in Figure S3.

**Figure 2:**
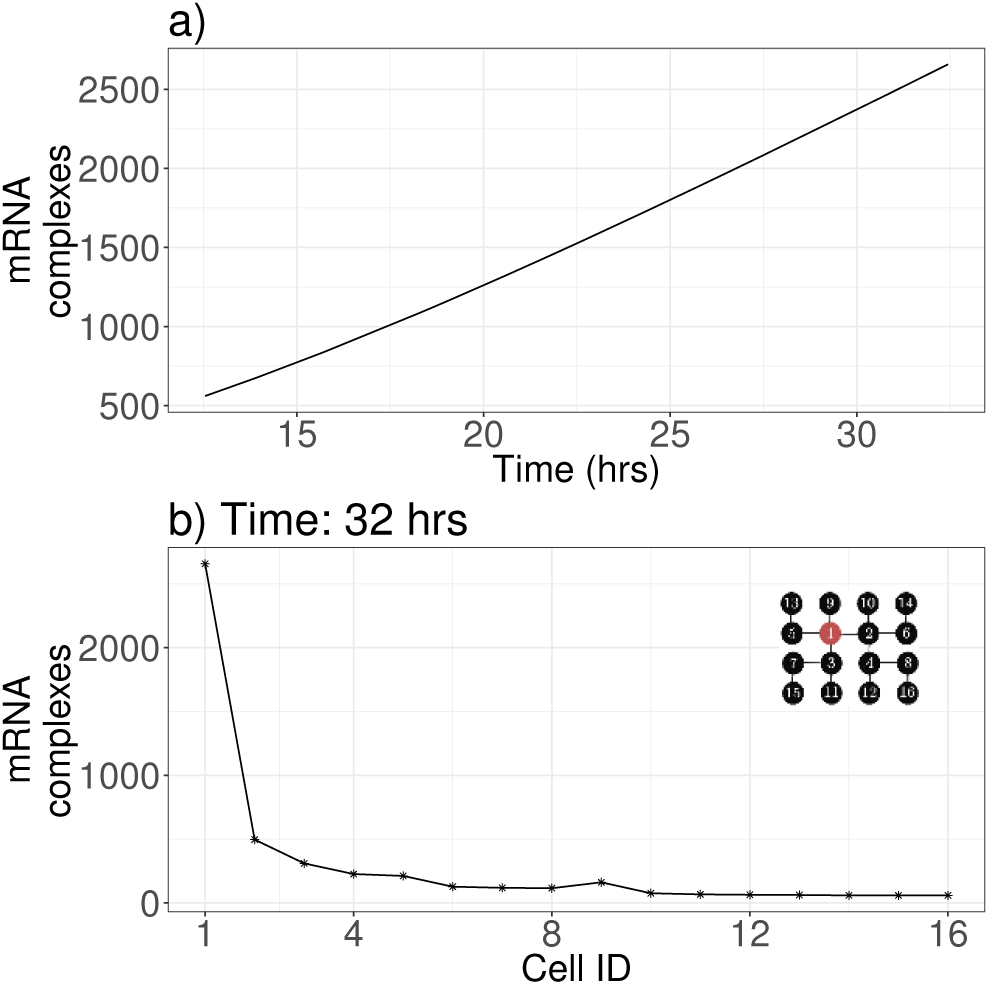
Predictions from the ODE model (1). (a) D behaviour of the number of complexes in the oocy remaining nurse cells are shown in Figure S3. (b) T tribution of complexes across the cells in the egg c at *t* = 32 hrs demonstrates the accumulation of in the oocyte. The numbering for the Cell ID on the *x* axis corresponds to that given in Figure 1. Parameters are *a* = 10 particles hr^*-*1^, *b* = 0.20 hr^*-*1^ and *ν* = 0.10.

### 3.2 Bayesian inference framework

We connect the model directly to quantitative experimental data so that it can provide relevant predictions and insight. We use a Bayesian inference framework to take account of both measurement and process uncertainty, incorporating the mechanistic model stated in Equation (1), a model of the measurement process, and prior knowledge of parameters.

The biological process model (Equation (1)) can be related to the observed data via a measurement model. The measurement model accounts for any errors in processing the data, in addition to raw experimental error. However, we first note that RNA complexes consist of multiple individual mRNA transcripts, and recall that upon entry to the oocyte there is some higher-order assembly of these complexes [12, 17, 18]. Since we have quantitative data on the number of RNA complexes in each cell, rather than the number of individual transcripts, we account for this higher-order assembly of complexes in the oocyte in the measurement model. We assume that in the oocyte we observe *ϕ y*_1_ complexes, where *ϕ* ∈ (0, 1] is a scaling parameter. We can interpret *ϕ* as a ratio between the number of transcripts per complex in the nurse cells as compared to the oocyte.

We use a negative binomial distribution for the measurement model, so that measured RNA complex counts, **z**, are given by

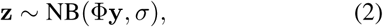

where Φ is a diagonal matrix with entries [*ϕ,* 1, 1, *…,* 1], *σ* is a parameter controlling the magnitude of the measurement error, and **y** is the solution of the biological process model (Equation (1)), giving the numbers of RNA complexes in each cell. Details on the parameterization for the negative binomial distribution are given in Supplementary Material Section E. We take the product of the likelihoods over each of the observed cells (including nurse cells and oocyte), with each biological sample corresponding to a unique time point, to give the full likelihood. Full details on the choice of prior distributions for parameters are given in Supplementary Material Section F. The parameters to be inferred are *a, b, ν, ϕ* and *σ*.

### 3.3 Quantification of higher-order assembly of gurken RNA complexes in the oocyte

Considering directly the total integrated intensity of RNA complexes in the oocyte compared to those in the nurse cells allows us to directly estimate the relative numbers of mRNA transcripts in an RNA complex in the oocyte compared to the nurse cells. We calculate the total integrated intensity for 448 foci in *n* = 13 example datasets for egg chambers from stage 5 to stage 8, including both the nurse cells and oocyte. By subtracting the background intensity, and dividing by the mean nurse cell intensity, we can obtain an estimate for *ϕ*, as shown in Figure 3: the median estimate gives *ϕ* = 0.345 *±* 0.048. Fitting a mixture of Gaussians model to the intensity data reveals how transcripts are packed within RNA complexes via multimodal distributions of intensities (Supplementary Figure S5).

**Figure 3:**
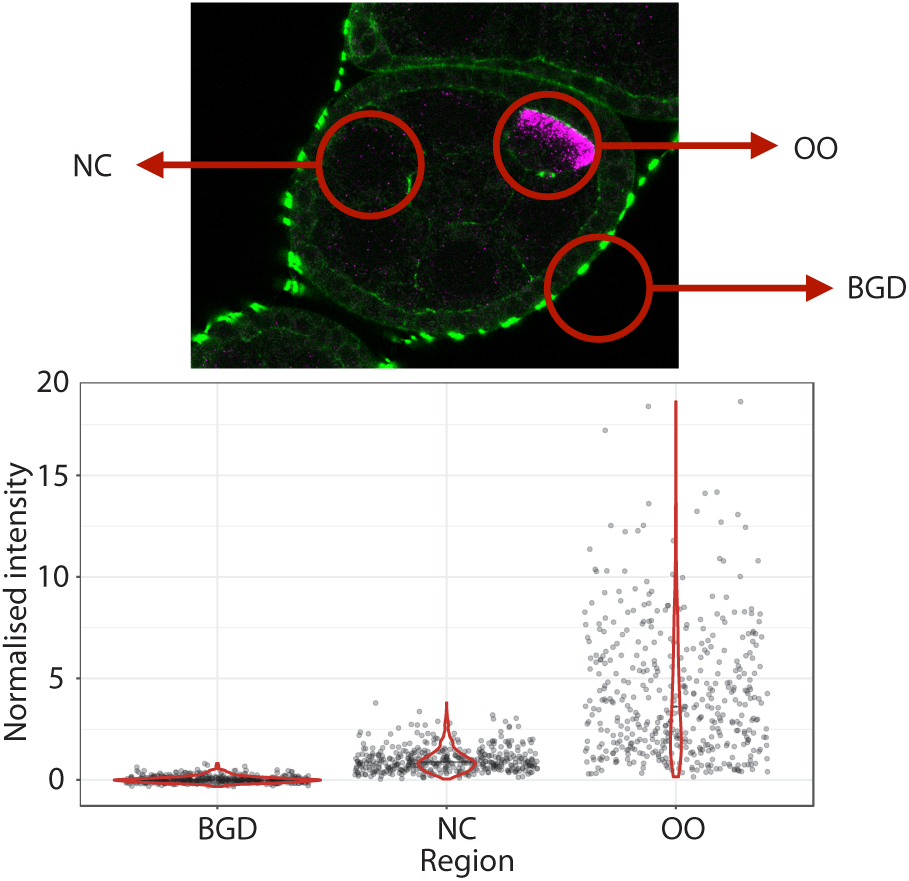
Assembly of higher order *gurken* complexes in the oocyte and quantification of the size of the assemblies in terms of equivalent complexes in the nurse cells. Typical complexes in background (BGD), nurse cells (NC) and the oocyte (OO) are identified from the circled regions indicated. By measuring total integrated intensity of these complexes in *n* = 13 egg chambers, we obtain distributions in each region, normalized by subtracting the average background value and scaling to a single nurse cell complex. We find that complexes in the oocyte are equivalent to a median of 2.5 times a single complex in the nurse cells.

**Figure 4:**
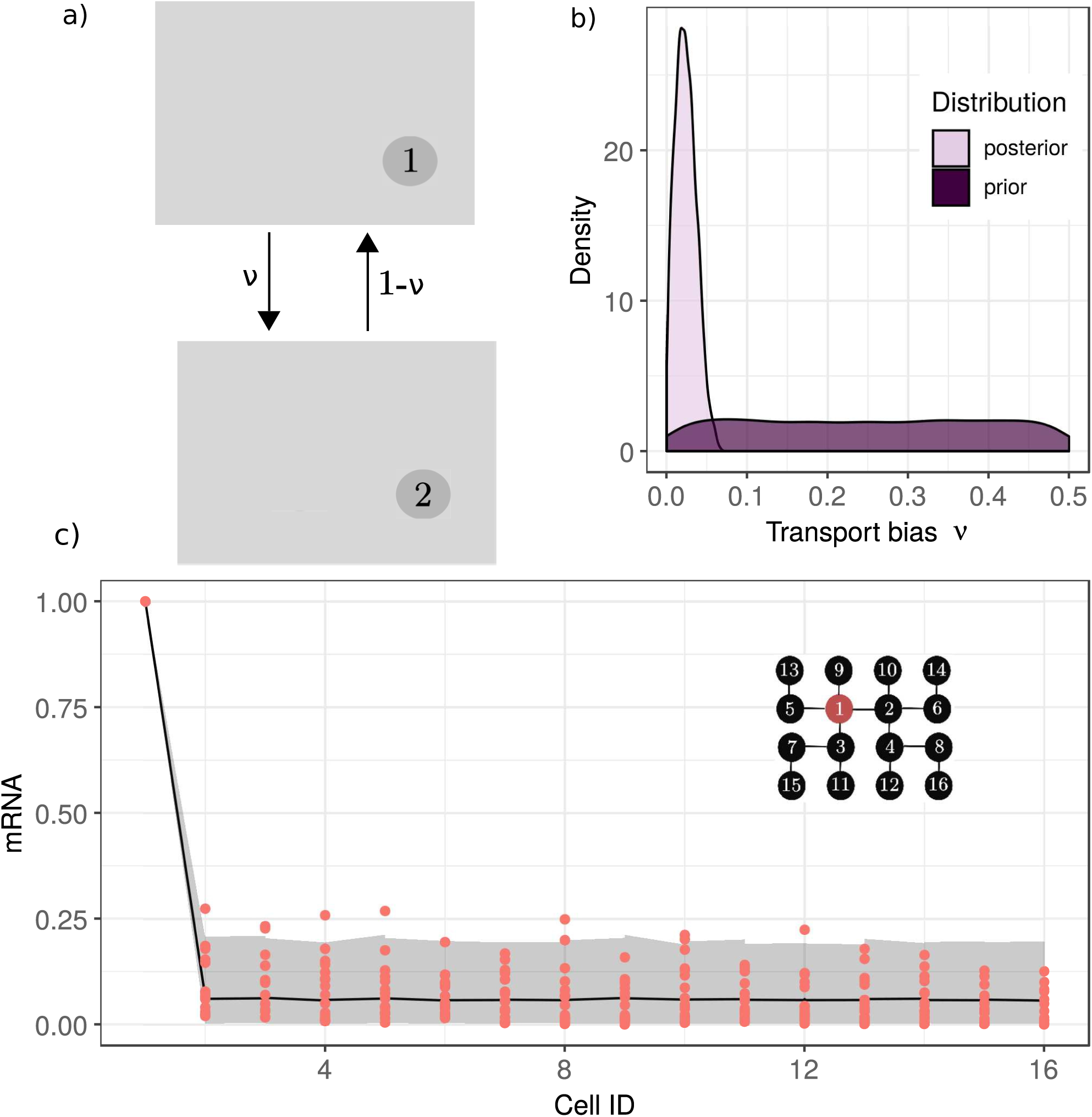
Results of MCMC sampling for the coarse grained model at steady state to determine the bias in transport through ring canals, as described in Section 3.4. In a), we display a schematic diagram of biased transport between two compartments, with transport from cell 2 to cell 1 at rate 1 − *ν*, and transport from cell 1 to cell 2 at rate *ν*. In b), we display a density plot of the marginal posterior distribution for the transport bias parameter *ν*, compared to the prior for the same parameter which is uniform over [0.0, 0.5]. This shows evidence of strong bias in transport through ring canals. In c), we show the posterior predictive distribution with the raw data shown as red points. The shaded region shows a 95% credible interval of the distribution of predictions from the model. Here the distribution of mRNA across cells in the egg chamber is normalized such that the amount of mRNA in the oocyte is 1. The model provides a good fit to the data, since most of the data points lie within the grey envelope of the 95% credible interval.

We can now use this independent estimate of *ϕ* to specify a strong prior for *ϕ* when fitting the full Bayesian model. We obtain posterior estimates for the assembly parameter, *ϕ*, with 0.40 as a median value, and [0.31, 0.49] as a 95% credible interval. This gives an estimate of between two and four nurse cell RNA complexes assembled into one equivalent oocyte RNA complex. The posterior distribution for *ϕ* is shown in pairwise plots of all the model parameters in Figure S6.

### 3.4 Biased transport through ring canals revealed by model comparison at quasi-steady state

We now consider the question of whether transport of RNA complexes through ring canals in *Drosophila* is unidirectional (towards the oocyte) or bidirectional (both towards and away from the oocyte). In addition, if transport is bidirectional, how biased is it towards the ooctye? Within the mathematical model (Section 3.1), the network structure (encoded in the matrix **B**) and the bias parameter, *ν*, impose a distribution of RNA complexes across the nurse cells. By inferring a posterior distribution for *ν*, through comparison of the model with quantitative data, we can estimate the extent to which transport through the ring canals is biased.

To simplify the model and ensuing Bayesian analysis, we consider the model behaviour in the large time, quasisteady-state limit (which corresponds to the time scale on which observations are made). In this limit, we have **y ≈** 15*a***k_2_***t*, where **k_2_** is the quasi-steady-state distribution that can be uniquely calculated for a given matrix **B** (see Supplementary Material Section H). Normalizing both the vector **k_2_** (so that the normalized RNA complex count in the oocyte is unity), and the equivalent distribution measured from smFISH data, enables comparison of the distribution of complexes across the egg chamber under the assumption of a Gaussian measurement error model, *N* (0, *ξ*^2^). We note that a different measurement error model is required here compared to Section 3.2, since the normalized mRNA distribution takes positive real values rather than integer counts. Priors are described in detail in Supplementary Material Section F.

In Figure 4, we show the marginal posterior distribution for the transport bias parameter, *ν*. The median of the marginal posterior distribution for *ν* is 0.02, with a 95% credible interval of [0.00, 0.05]. The posterior predictive distribution (Figure 4(c)) contains the observed data points within the 95% credible region for the complex distribution in each cell. Together these results strongly support the hypothesis that transport of complexes is significantly biased towards the oocyte.

### 3.5 Estimates of production and transport rates show production and transport are in tightly regulated balance

We obtain estimates for the rates of production, *a*, and transport, *b*, of RNA complexes by applying the modelling approach outlined in Equation (1) and Section 3.2. We consider the full dynamic behaviour of the model, together with the smFISH dataset of *n* = 16 egg chambers (see Section 2) and infer all model parameters (*a, b, ν, ϕ* and *σ*).

Upon sampling from the posterior for the full model, we find that the production rate, *a*, takes a median value of 10.0 and lies within a 95% credible interval of [7.7, 14.1] (in units of [particles hr^−1^]). The transport rate, *b*, has a median value of 0.22 and lies within a 95% credible interval of [0.16, 0.31] (in units of [hr^−1^]). The marginal posterior distributions for the rate parameters *a* and *b* are shown in Figure 5, and the posterior distribution for all the model parameters is shown pairwise in Figure S6.

**Figure 5:**
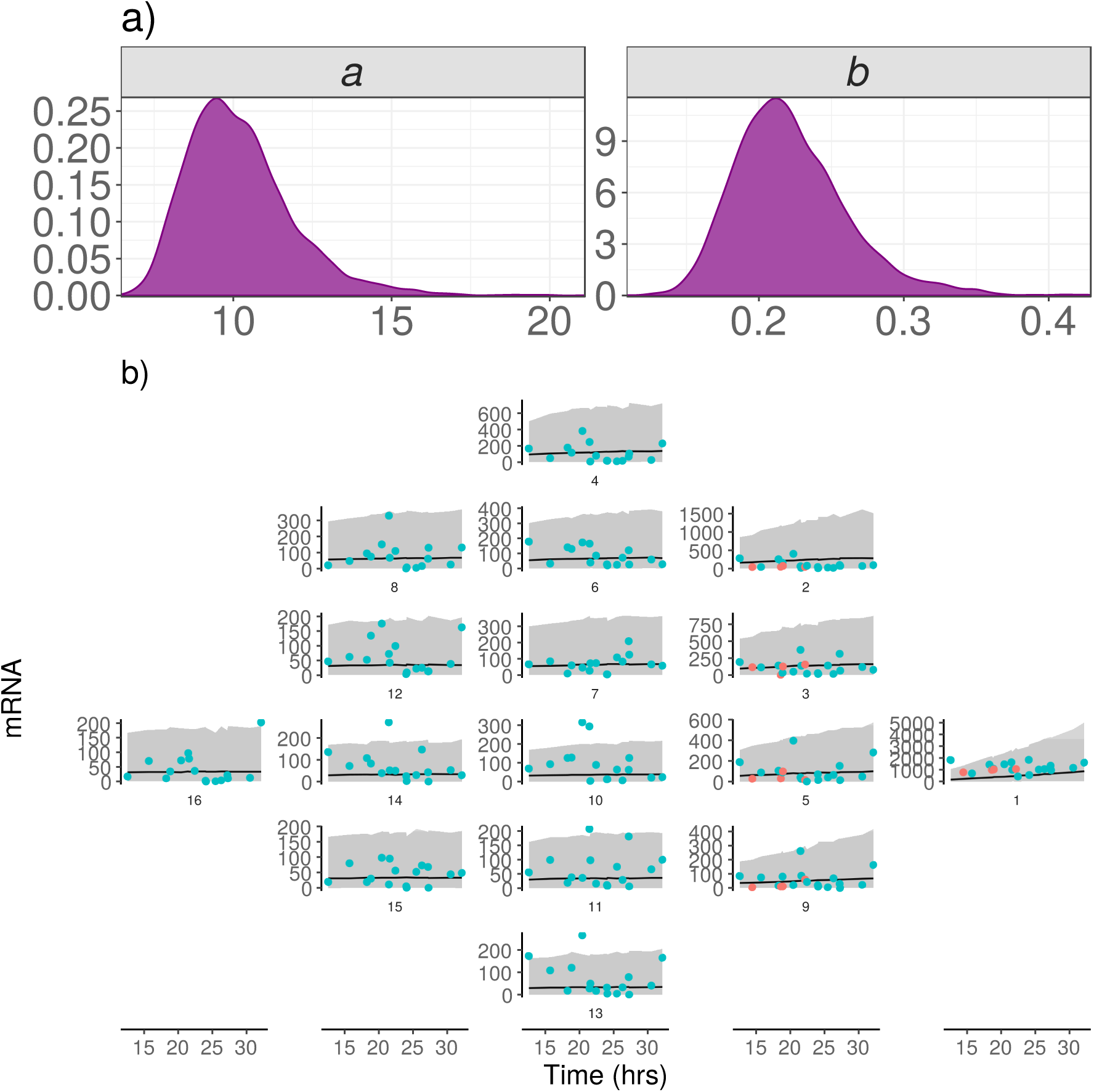
Results of MCMC sampling for the model described in Section 3.2 and sensitivity to model parameters. In a), we display the posterior marginal distribution for the rate parameters *a* and *b*. In b), we show the posterior predictive distribution, which gives the predictions from the model with parameters fitted to the data. Each panel gives the prediction for each cell (numbered 1 for the oocyte through 2 to 16 for the nurse cells as in Figure 1) as a solid line, with measured experimental particle counts shown as points. A 95% credible interval for the predictions is shown as a shaded region. The blue points are observed data used to train the model, whilst the red points are observed data used to test the model (note that these red points are egg chambers for which only the oocyte and neighbouring nurse cells have been segmented). The subplots are arranged to highlight distance from the oocyte within the network of connections between nurse cells shown in Figure 1c).

Using these parameter estimates, we can compare the relative contributions of the production and transport terms in Equation (1) to the rate of change in the number of RNA complexes. We have 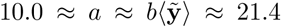, where *a* is the complex production rate, *b* is the transport rate and 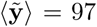 is the median number of *gurken* RNA complexes at time *t* considered across all cells of the egg chamber. This result indicates that the RNA complex production rate is of the same order of magnitude as the rate of complex transport, and therefore suggests that production and transport are tightly balanced. We provide further evidence to support this hypothesis by evaluating the sensitivity of the model described in Section 3.1 to changes in the rate parameters *a* and *b* (see Supplementary Material Section J).

### 3.6 Perturbing mRNA production yields results inconsistent with model predictions

To further validate our results, we explored whether the model can predict the response of the system to a perturbation in the RNA complex production rate by considering an over-expression mutant with multiple copies of the gurken gene. We make the assumption that the mutant has RNA complex production rate *γa*, where *a* is the production rate in wild type, and *γ >* 1 is a scale factor, but that all other model parameters are unchanged.

The model specified in Equation (1) predicts that the total number of RNA complexes in an egg chamber should increase at rate 15*a* in wild type, and 15*γa* in the over-expression mutant. Comparing this prediction with experimental data, we estimate a median value for *γ* of 2.23, with a 95% credible interval of [1.38, 3.06] (Supplementary Material Section L). We take the value *γ* = 2.23 for the remainder of this work.

We now use the model to predict the distribution of RNA complexes in each of the cells of the egg chamber by sampling from the posterior predictive distribution generated in Section 3.5 using wild type data only. In doing so, the implicit assumption is that, in the over-expression mutant, the only process significantly affected is the RNA complex production rate. For the oocyte and neighbouring nurse cells, the model predictions are consistent with the observed data (Figure 6). However, for the nurse cells furthest away from the oocyte (such as 8, 12, 14, 15, 16 which are at least three ‘steps’ away from the oocyte) the observed experimental data contains nurse cells with much greater accumulation of RNA complexes than predicted by the model (Figure 6). We highlight further the phenotypic differences in the distribution of mRNA across cells based on distance from the oocyte in Figure S9.

**Figure 6:**
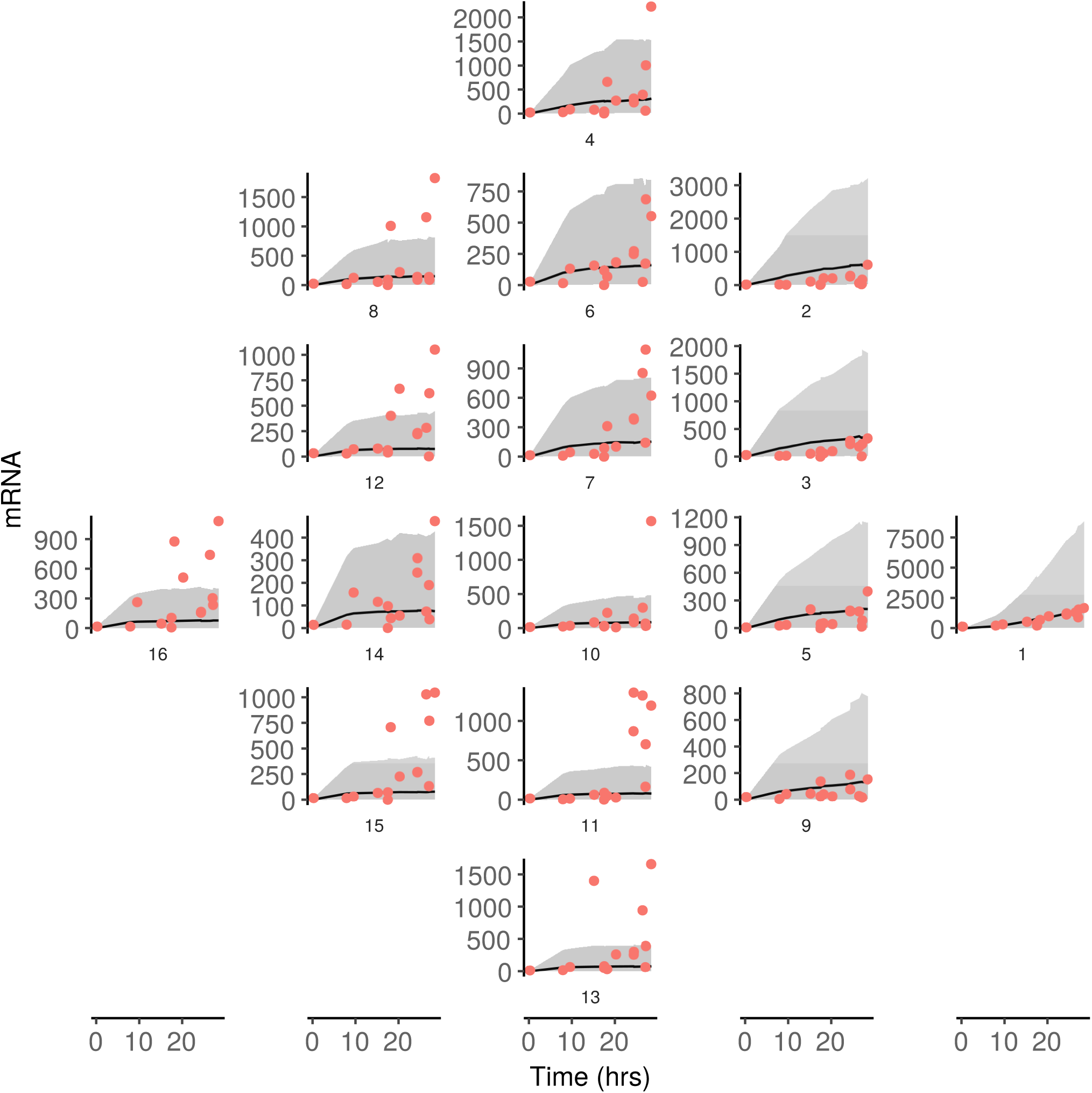
Posterior predictive distribution for the over-expression mutant based on the ODE model (1) fitted to wild type data. Model predictions (gray envelope) show good agreement with observed data (red points) for cells close to the oocyte. Cells further from the oocyte, such as cell 16, show discrepancy between model predictions and the observed data, indicating there are biological mechanisms not captured by the model. The observed data for the over-expression mutant is shown as red dots, with the gray envelope showing a 95% credible interval of predictions from the model via the posterior predictive distribution. The subplots are arranged to highlight distance from the oocyte within the network of connections between nurse cells shown in Figure 1c).

Crucially, the lack of agreement between predictions and observed data clearly indicates that the model does not capture key mechanistic details of the mRNA localization process that are relevant to the over-expression mutant. The aim of the ensuing analysis is to tease apart the nature of these missing mechanistic details.

### 3.7 Localization of RNA in the oocyte of the over-expression mutant reveals robustness

Quasi-steady-state analysis of the ODE model in Equation (1) predicts that RNA complex numbers in the oocyte, *y*1, grow linearly at a rate proportional to 15*a* (Supplementary Material Section H). Therefore, in addition, the model predicts that in the over-expression mutant, which has an increased production rate of *γa*, the number of RNA complexes localized to the oocyte at a given developmental time should increase by a factor of *γ* compared to wild type. However, rather than this predicted increase, we observe the number of RNA complexes in the over-expression mutant oocytes to be similar to that in wild type (Figure 7). This results demonstrates an incredible robustness of the system in the sense that the same amount of *gurken* RNA is localized to the oocyte, despite large changes to the production rate. A key question is now what mechanism gives rise to the observed robustness?

**Figure 7:**
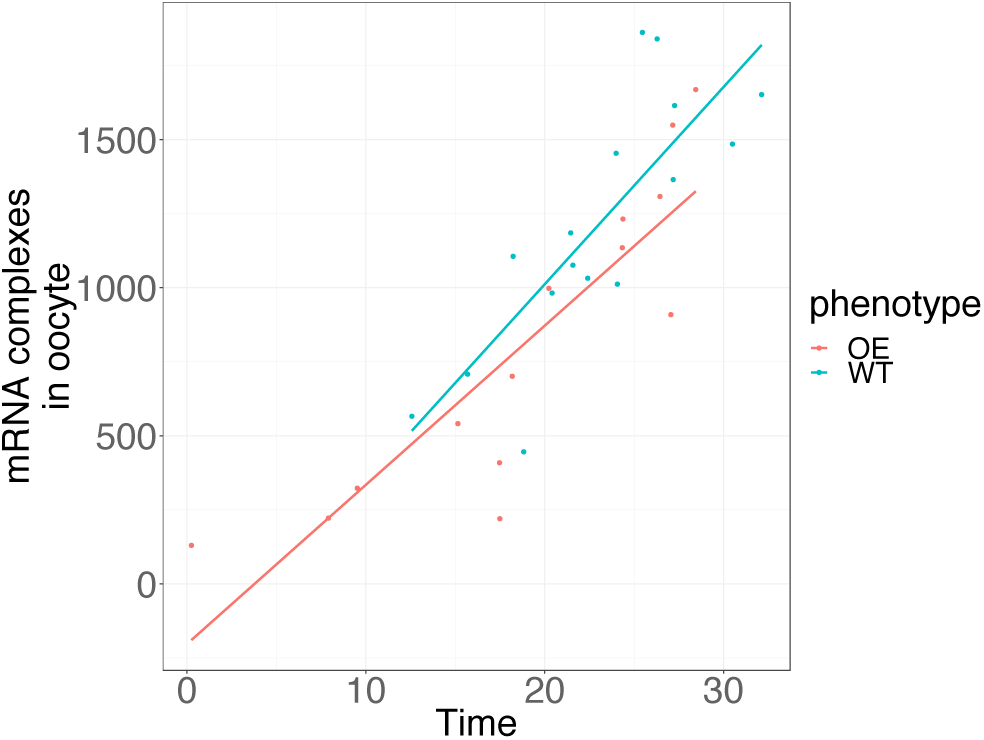
The number of mRNA complexes in the oocyte increases linearly over time for both the wild type (blue) and the over-expression mutant (red). Strikingly, the rate of increase of mRNA in the oocyte (shown by the gradient of the red and blue lines) is approximately equal in each case, whereas we would have expected this to be *γ* times greater for the over-expression mutant. The tight regulation of the localization process biologically ensures the right amount of RNA reaches the oocyte at the right time, even under biological perturbation. Here this is due to blocking of ring canals. The points show observed data on number of RNA complexes, and the lines show linear models fit to these data.

### 3.8 Biological mechanisms to explain the over-expression data can be represented as alternative models

We propose several plausible biological mechanisms to explain the behaviour of the localization process in the over-expression mutant, as listed below, and evaluate their ability to recapitulate data from the over-expression mutant.

#### Inhomogeneous production

Previously we assumed equal production rates of RNA complexes across all nurse cells. However, the over-expression of *gurken* in nurse cells is driven by the GAL4-UAS system, which can result in patchy expression across cells. We now relax the assumption of equal production rates, and estimate the production rates of individual nurse cells by quantifying the number of nascent *gurken* transcripts (Supplementary Material N).

#### Blocking or queuing at ring canals

With increased levels of *gurken* in the over-expression mutant, the environment inside the nurse cells is more crowded. We hypothesize that the transport of RNA complexes through ring canals could be blocked or restricted due to crowding. Blocking in this way restricts transport both towards, and away from, the oocyte. We can represent this hypothesis in the model by changing appropriate entries in the matrix **B** to zero: manual examination of the data was used to predict which ring canals are blocked.

#### Density dependent transport

The final hypothesis we consider is that increases in *gurken* in the over-expression mutant lead to saturation of the transport mechanism. This hypothesis is motivated by observations of a build up of RNA at nuclear pore complexes in the over-expression mutant, suggesting that nuclear export or the ability to form competent mRNA particles for transport may be saturated. Instead of assuming that transport rate between cells is linearly proportional to the number of RNA complexes, we assume a saturating transport rate of the form

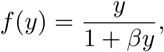

where *β* is a parameter describing the density dependence.

We represent the hypotheses detailed above as a collection of models, ℳ = {M0, M1, …, M7}, that together represent all possible combinations of the additional mechanisms (see Table 1 for details). Models including the crowding-induced blocking at ring canals and inhomogeneous production mechanisms can be forward simulated using parameters from the posterior distribution based on wild type data, as in Figure 6. Models including density dependent transport must be fitted to the wild type data to estimate the parameter *β*.

**Table 1:** A description of the collection of models ℳ = {M0, M1, …, M7}, together with the weights generated using model comparison (see Section 2.7 for details).

|  | Production | Blocking | Density dependent transport | Pseudo BMA+ weights | Stacking weights |
| --- | --- | --- | --- | --- | --- |
| M0 | 0 | 0 | 0 | 0.04 | 0.00 |
| M1 | 0 | 1 | 0 | 0.94 | 0.40 |
| M2 | 0 | 0 | 1 | 0.00 | 0.08 |
| M3 | 1 | 0 | 0 | 0.01 | 0.23 |
| M4 | 1 | 0 | 1 | 0.00 | 0.00 |
| M5 | 1 | 1 | 0 | 0.00 | 0.29 |
| M6 | 0 | 1 | 1 | 0.00 | 0.00 |
| M7 | 1 | 1 | 1 | 0.00 | 0.00 |

### 3.9 Model comparison reveals blocking of ring canals as the most plausible mechanism

We use model comparison approaches (Section 2.7) to evaluate the ability of the different models to explain the biological observations. Table 1 indicates strong support for the blocking of ring canals as a mechanism to explain the observed over-expression data: Model M1 has a pseudo-BMA+ weight close to 1, indicating that of the models considered this is the most plausible model. The more conservative stacking weights approach also weights several other models relatively highly, however these are predominantly models that involve blocking as well as additional mechanisms. These results suggest that none of the models considered perfectly describe the data generating process (to be expected since the model is an incredibly simplified description of a very complex biological process). However, predictions of RNA complex distributions for the over-expression mutant made using M1 (which incorporates only the blocking of ring canals) are substantially improved compared to the predictions of the basic model. Taken together, our results reveal crowding-driven blocking of ring canals as the most plausible explanation for our experimental observations.

### 3.10 Reexamining microscopy data provides evidence of queuing at ring canals

A natural question to ask is then whether it is possible to validate this prediction experimentally? By re-examining microscopy data from the over-expression mutant, we demonstrate evidence for the crowding-related blocking of ring canals in smFISH data: we observe clusters of RNA complexes around ring canals (Figure 8). Similar observations of a build up of RNA complexes near ring canals have also been reported upon injection of *gurken* mRNA in nurse cells [12]. Together, these results suggest that, while the transport of RNA complexes within nurse cells may be relatively fast, the efficiency of the transport process as a whole can suffer when transit through ring canals is blocked by crowding. Based on assessment of the mRNA counts across cells (Supplementary Figure S12), we find 64% (*n* = 14) of the over-expression mutant egg chambers show evidence of blocking behaviour. The mechanism leading to crowding-induced blocking in over-expression mutants may be present also in wild type, but such events occur rarely in wild type due to lower accumulation of RNA complexes within a nurse cell.

**Figure 8:**
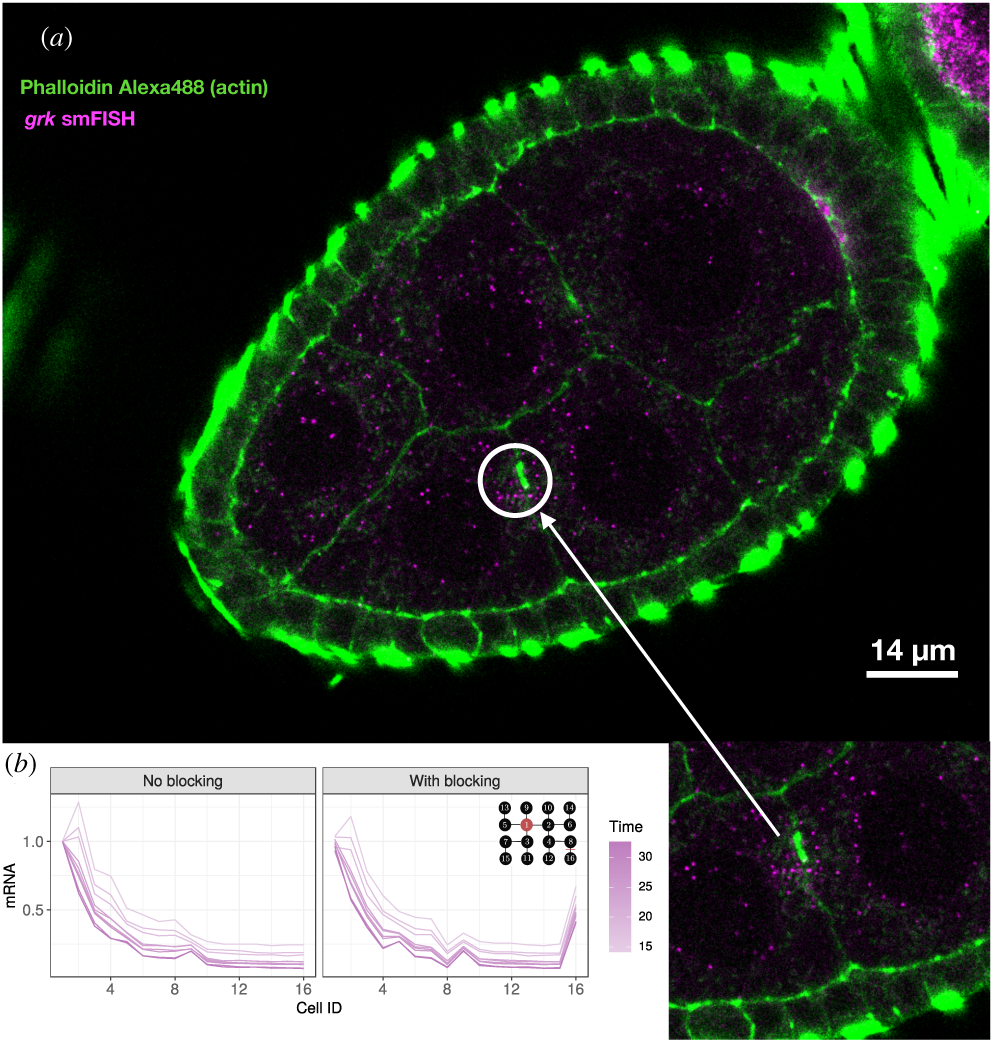
Blocking of ring canals may result in significant accumulation of mRNA in nurse cells far from the oocyte. a) A clustered distribution of mRNA complexes queuing near the ring canal (arrow). b) A comparison between simulations from the model with all ring canals transporting RNA equally, and a situation where the ring canal between cells 8 and 16 is blocked. The schematic inset in a) shows the position of the blocked ring canal within the tissue.

## 4 Discussion

Combining simple mechanistic mathematical models and Bayesian inference approaches together with experimental evidence, we have investigated the dynamics of mRNA localization in *Drosophila* egg chambers, and made quantitative predictions of the dynamics of the localization process. Estimates of the production and transport rates of *gurken* mRNA indicate that these processes are tightly balanced in mRNA localization. To confer robustness on the system, large perturbations must be buffered by the localization mechanism to ensure that the ‘correct’ amount of RNA is localized at the right time in development. We have shown, using a simple model, that crowding-driven blocking of ring canals is sufficient to regulate the localization of mRNA in the oocyte.

In addition, our results suggest that transport of RNA complexes through ring canals is strongly biased. The characteristic network of connections between the nurse cells specifies a given network structure that we represent via the matrix **B** in Equation (1). By inferring the bias parameter, *ν*, we have shown that the observed distribution of RNA complexes across the egg chamber is consistent with strongly biased transport of RNA complexes towards the oocyte. This strong bias at a macroscopic cellular scale is nonetheless consistent with weak bias in transport at a microscopic scale on the microtubule network [51], since averaging over many movements of many particles moving with weakly biased random motion can result in strong bias for the ensemble average.

We have also provided measurements and estimates for the assembly of higher order RNA complexes in the oocyte following remodelling upon transit through the ring canals between the oocyte and the nurse cells. Measurements of RNA content for *bicoid, oskar* and *nanos* RNA complexes have been performed previously [18, 52], although largely for later stages of development, and we note that the estimates given here are consistent with similar measurements on the distribution of multimeric RNA complexes of *oskar* upon entry to the oocyte at stage 10b [18]. This process of remodelling of RNA complexes upon transit into the oocyte may play a role within wider changes in how RNA is packaged in the oocyte compared to the nurse cells.

We attempted to validate the mathematical model (Equation (1)) by making predictions about behaviour of an over-expression mutant. The predictions of this simple model revealed a discrepancy in the accumulation of mRNA in the nurse cells furthest from the oocyte compared with the observed experimental data: in some cases, far more mRNA is observed in these cells than predicted by the model. In addition, the observed rate of localization of mRNA in the oocyte is similar in the over-expression mutant as compared to wild type, contrary to predictions from the simple model. We resolved this discrepancy by using model comparison approaches to compare the ability of a range of biological mechanisms to replicate the observed data. Our results suggest crowding-induced blocking of ring canals as a plausible explanation for the discrepancy between model prediction and experimental observations, and more generally that the transport process becomes overloaded in the over-expression mutant. We therefore argue that the crowding-induced blocking of ring canals is effectively a regulatory mechanism that helps ensure the robustness of *gurken* localization in *Drosophila* oocytes. Precise control of gene expression during development can be facilitated by indirect methods [53]; blocking of ring canals is an example of such indirect regulation.

We have considered generalizations of the simple model (Equation (1)) to incorporate inhomogeneous variance across cells in our observations or a decay term with decay of RNA at rate *δ*. Details are described in Supplementary Material Sections P and Q. Our conclusions about the blocking mechanism hold also for these more general descriptions. The simple model is favoured by model comparison when compared to these models, which do not generalize as well to predictions for the over-expression data. Alternative hypotheses to account for the discrepency between model predictions and observations, based on different degradation rates for the different phenotypes, may have some merit, but they are hard to assess within the framework of prediction conditioned on wild type data used to compare the other candidate models (see Supplementary Material Section P).

To further assess the crowding-induced blocking mechanism, we suggest directly measuring transit through ring canals in the wild type and over-expression mutant conditions. However, this task is nontrivial and would require extended time lapse imaging to track movements of complexes through representative subsets of all ring canals.

Finally, we highlight that the conclusions drawn as a result of this combined modelling-experiment study were made possible through use of Bayesian inference approaches that allowed us to interrogate and interpret quantitative data, making full use of the information contained therein. It is important to note also that our conclusions were reached through the use of an incredibly simple, analytically tractable, coarse-grained model, that allowed us to abstract many of the intricate details, and focus on salient mechanisms. We hope that our work serves as an exemplar for future studies in this area.

## Author contributions

JUH and RMP designed and performed biological experiments and JUH carried out primary analysis and interpretation of the data. JUH and REB developed the modelling approaches and JUH wrote the code and performed statistical inference. All authors discussed the results and conclusions, commented on and contributed to the revision of the manuscript. REB, RMP and ID supervised the project.

## Acknowledgements

This work was supported by funding from the Engineering and Physical Sciences Research Council (EPSRC) (grant no. EP/G03706X/1). R.E.B. is a Royal Society Wolfson Research Merit Award holder and a Leverhulme Research Fellow, and also acknowledges the BBSRC for funding via grant no. BB/R000816/1. This work was additionally supported by a Wellcome Trust Senior Research Fellowship (096144) and Wellcome Investigator Award (209412) to I.D. and supporting R.M.P. Thanks also to MICRON (http://micronoxford.com, supported by Wellcome Strategic Awards 091911/B/10/Z and 107457/Z/15/Z) for access to equipment.

## Footnotes

1 In referring to production, we include mRNA transcription, export through the nuclear pore complex, and assembly of RNA-protein complexes.

## References

[1] Becalska, A. N., and E. R. Gavis, 2009. Lighting up mRNA localization in Drosophila oogenesis. Development 136:2493–2503.

[2] Buxbaum, A. R., G. Haimovich, and R. H. Singer, 2014. In the right place at the right time: visualizing and understanding mRNA localization. Nature Reviews Molecular Cell Biology.

[3] Parton, R. M., A. Davidson, I. Davis, and T. T. Weil, 2014. Subcellular mRNA localisation at a glance. Journal of Cell Science 127:2127–2133.

[4] Wolpert, L., R. Beddington, J. Brockes, T. Jessell, P. Lawrence, and E. Meyerowitz, 1998. Principles of Development. Oxford University Press, Oxford.

[5] Wilkie, G. S., and I. Davis, 2001. Drosophila wingless and pair-rule transcripts localize apically by dynein-mediated transport of RNA particles. Cell 105:209–219.

[6] Bobola, N., R. Jansen, T. H. Shin, and K. Nasmyth, 1996. Asymmetric accumulation of Ash1p in postanaphase nuclei depends on a myosin and restricts yeast mating-type switching to mother cells. Cell 84:699–709.

[7] Mowry, K. L., and D. A. Melton, 1992. Vegetal messenger RNA localization directed by a 340-nt RNA sequence element in Xenopus oocytes. Science 255:991–994.

[8] Rosbash, M., and R. H. Singer, 1993. RNA travel: tracks from DNA to cytoplasm. Cell 75:399–401.

[9] Nevo-Dinur, K., A. Nussbaum-Shochat, S. Ben-Yehuda, and O. Amster-Choder, 2011. Translation-independent localization of mRNA in E. coli. Science 331:1081–1084.

[10] Cáceres, L., and L. A. Nilson, 2005. Production of gurken in the nurse cells is sufficient for axis determination in the Drosophila oocyte. Development 132:2345–2353.

[11] Bullock, S. L., and D. Ish-Horowicz, 2001. Conserved signals and machinery for RNA transport in Drosophila oogenesis and embryogenesis. Nature 414:611–616.

[12] Clark, A., C. Meignin, and I. Davis, 2007. A Dynein-dependent shortcut rapidly delivers axis determination transcripts into the Drosophila oocyte. Development 134:1955– 1965.

[13] Spradling, A. C., 1993. Developmental genetics of oogenesis. In The development of Drosophila melongaster. Cold Spring Harbour, NY, Cold Spring Harbour Laboratory Press, 1–70.

[14] Alsous, J. I., P. Villoutreix, N. Stoop, S. Y. Shvartsman, and J. Dunkel, 2018. Entropic effects in cell lineage tree packings. Nature Physics 14:1016.

[15] Femino, A. M., F. S. Fay, K. Fogarty, and R. H. Singer, 1998. Visualization of single RNA transcripts in situ. Science 280:585–590.

[16] Raj, A., P. Van Den Bogaard, S. A. Rifkin, A. Van Oudenaarden, and S. Tyagi, 2008. Imaging individual mRNA molecules using multiple singly labeled probes. Nature Methods 5:877–879.

[17] Bohrmann, J., and K. Biber, 1994. Cytoskeleton-dependent transport of cytoplasmic particles in previtellogenic to mid-vitellogenic ovarian follicles of Drosophila: time-lapse analysis using video-enhanced contrast microscopy. Journal of Cell Science 107:849–858.

[18] Little, S. C., K. S. Sinsimer, J. J. Lee, E. F. Wieschaus, and E. R. Gavis, 2015. Independent and coordinate trafficking of single Drosophila germ plasm mRNAs. Nature Cell Biology 17:558.

[19] Trong, P. K., H. Doerflinger, J. Dunkel, D. St Johnston, and R. E. Goldstein, 2015. Cortical microtubule nucleation can organise the cytoskeleton of Drosophila oocytes to define the anteroposterior axis. eLife 4:e06088.

[20] Ciocanel, M. V., J. A. Kreiling, J. A. Gagnon, K. L. Mowry, and B. Sandstede, 2017. Analysis of Active Transport by Fluorescence Recovery after Photobleaching. Biophysical Journal 112:1714–1725.

[21] Bressloff, P. C., and J. M. Newby, 2013. Stochastic models of intracellular transport. Reviews of Modern Physics 85:135.

[22] Howard, J., 2001. Mechanics of Motor Proteins and the Cytoskeleton. Sinauer, Sunderland, MA.

[23] Hafner, A. E., and H. Rieger, 2018. Spatial cytoskeleton organization supports targeted intracellular transport. Biophysical Journal 114:1420–1432.

[24] Hawkins, R. J., O. Benichou, M. Piel, and R. Voituriez, 2009. Rebuilding cytoskeleton roads: Active-transport-induced polarization of cells. Physical Review E 80:040903.

[25] Bressloff, P. C., and B. Xu, 2015. Stochastic active-transport model of cell polarization. SIAM Journal on Applied Mathematics 75:652–678.

[26] Alsous, J. I., P. Villoutreix, A. M. Berezhkovskii, and S. Y. Shvartsman, 2017. Collective growth in a small cell network. Current Biology 27:2670–2676.e4.

[27] Bökel, C., S. Dass, M. Wilsch-Bräuninger, and S. Roth, 2006. *Drosophila* Cornichon acts as cargo receptor for ER export of the TGF*α*-like growth factor Gurken. Development 133:459–470.

[28] Jaramillo, A. M., T. T. Weil, J. Goodhouse, E. R. Gavis, and T. Schupbach, 2008. The dynamics of fluorescently labeled endogenous gurken mRNA in Drosophila. Journal of Cell Science 121:887–894.

[29] Weil, T. T., R. M. Parton, B. Herpers, J. Soetaert, T. Veenendaal, D. Xanthakis, I. M. Dobbie, J. M. Halstead, R. Hayashi, C. Rabouille, and I. Davis, 2012. Drosophila patterning is established by differential association of mRNAs with P bodies. Nature Cell Biology 14:1305–1313.

[30] Edwards, K. A., M. Demsky, R. A. Montague, N. Weymouth, and D. P. Kiehart, 1997. GFP-Moesin illuminates Actin Biophysical Journal 00(00) 1–14 cytoskeleton dynamics in living tissue and demonstrates cell shape changes during morphogenesis in Drosophila. Developmental Biology 191:103–117.

[31] Parton, R. M., A. M. Vallés, I. M. Dobbie, and I. Davis, 2010. Pushing the limits of live cell imaging in *Drosophila*. Live cell imaging: a laboratory manual 387–418.

[32] Davidson, A., R. M. Parton, C. Rabouille, T. T. Weil, and I. Davis, 2016. Localized translation of gurken/TGF-*α* mRNA during axis specification is controlled by access to Orb/CPEB on processing bodies. Cell Reports 14:2451–2462.

[33] Schindelin, J., I. Arganda-Carreras, E. Frise, V. Kaynig, M. Longair, T. Pietzsch, S. Preibisch, C. Rueden, S. Saalfeld, and B. Schmid, 2012. Fiji: an open-source platform for biological-image analysis. Nature Methods 9:676.

[34] Allan, C., J. M. Burel, J. Moore, C. Blackburn, M. Linkert, S. Loynton, D. MacDonald, W. J. Moore, C. Neves, and A. Patterson, 2012. OMERO: flexible, model-driven data management for experimental biology. Nature Methods 9:245.

[35] Linkert, M., C. T. Rueden, C. Allan, J. M. Burel, W. Moore, A. Patterson, B. Loranger, J. Moore, C. Neves, and D. MacDonald, 2010. Metadata matters: access to image data in the real world. The Journal of Cell Biology 189:777–782.

[36] Mueller, F., A. Senecal, K. Tantale, H. Marie-Nelly, N. Ly, O. Collin, E. Basyuk, E. Bertrand, X. Darzacq, and C. Zimmer, 2013. FISH-quant: automatic counting of transcripts in 3D FISH images. Nature Methods 10:277–278.

[37] Hailstone, M., L. Yang, D. Waithe, T. J. Samuels, Y. Arava, T. Dobrzycki, R. M. Parton, and I. Davis, 2017. Brain development: machine learning analysis of individual stem cells in live 3D tissue. bioRxiv.

[38] Yang, L., J. Titlow, D. Ennis, C. Smith, J. Mitchell, F. L. Young, S. Waddell, D. Ish-Horowicz, and I. Davis, 2017. Single molecule fluorescence in situ hybridisation for quantitating post-transcriptional regulation in *Drosophila* brains. bioRxiv 128785.

[39] Shimada, Y., K. M. Burn, R. Niwa, and L. Cooley, 2011. Reversible response of protein localization and microtubule organization to nutrient stress during Drosophila early oogenesis. Developmental Biology 355:250–262.

[40] Jia, D., Q. Xu, Q. Xie, W. Mio, and W. M. Deng, 2016. Automatic stage identification of *Drosophila* egg chamber based on DAPI images. Scientific Reports 6:18850.

[41] Carpenter, B., A. Gelman, M. Hoffman, D. Lee, B. Goodrich, M. Betancourt, M. Brubaker, J. Guo, P. Li, and A. Riddell, 2017. Stan: A Probabilistic Programming Language. Journal of Statistical Software 76:1–32.

[42] Betancourt, M., S. Byrne, S. Livingstone, and M. Girolami, 2017. The geometric foundations of Hamiltonian Monte Carlo. Bernoulli 23:2257–2298.

[43] Piironen, J., and A. Vehtari, 2017. Comparison of Bayesian predictive methods for model selection. Statistics and Computing 27:711–735.

[44] Vehtari, A., A. Gelman, and J. Gabry, 2017. Practical Bayesian model evaluation using leave-one-out cross-validation and WAIC. Statistics and Computing 27:1413– 1432.

[45] Vehtari, A., J. Gabry, Y. Yao, and A. Gelman, 2018. loo: Efficient leave-one-out cross-validation and WAIC for Bayesian models. https://CRAN.R-project.org/package=loo, rpackageversion2.0.0.

[46] Yao, Y., A. Vehtari, D. Simpson, and A. Gelman, 2018. Using stacking to average Bayesian predictive distributions. Bayesian Analysis.

[47] Tadros, W., and H. D. Lipshitz, 2005. Setting the stage for development: mRNA translation and stability during oocyte maturation and egg activation in *Drosophila*. Developmental Dynamics 232:593–608.

[48] Little, S. C., G. Tkačik, T. B. Kneeland, E. F. Wieschaus, and T. Gregor, 2011. The formation of the Bicoid morphogen gradient requires protein movement from anteriorly localized mRNA. PLOS Biology 9:e1000596.

[49] King, R. C., and R. G. Burnett, 1959. Autoradiographic study of uptake of tritiated glycine, thymidine, and uridine by fruit fly ovaries. Science 129:1674–1675.

[50] Schäling, B., 2011. The boost C++ libraries. Boris Schäling.

[51] Parton, R. M., R. S. Hamilton, G. Ball, L. Yang, C. F. Cullen, W. Lu, H. Ohkura, and I. Davis, 2011. A PAR-1–dependent orientation gradient of dynamic microtubules directs posterior cargo transport in the *Drosophila* oocyte. The Journal of Cell Biology 194:121–135.

[52] Trovisco, V., K. Belaya, D. Nashchekin, U. Irion, G. Sirinakis, R. Butler, J. J. Lee, E. R. Gavis, and D. St Johnston, 2016. bicoid mRNA localises to the *Drosophila* oocyte anterior by random Dynein-mediated transport and anchoring. eLife 5.

[53] Little, S. C., M. Tikhonov, and T. Gregor, 2013. Precise developmental gene expression arises from globally stochastic transcriptional activity. Cell 154:789–800.

