## Supplementary Material for "Testing models of mRNA localization reveals robustness regulated by reducing transport between cells"

#### A Connectivity matrix

Here we give the full connectivity matrix between nurse cells and the oocyte based on the characteristic connections due to cell divisions, with dependence on the transport bias parameter, $\nu$.

$$B = \begin{pmatrix}
-4\nu & 1-\nu & 1-\nu & 0 & 1-\nu & 0 & 0 & 0 & 1-\nu & 0 & 0 & 0 & 0 & 0 & 0 & 0 \\
\nu & -1-2\nu & 0 & 1-\nu & 0 & 1-\nu & 0 & 0 & 0 & 1-\nu & 0 & 0 & 0 & 0 & 0 & 0 \\
\nu & 0 & -1-\nu & 0 & 0 & 0 & 1-\nu & 0 & 0 & 0 & 1-\nu & 0 & 0 & 0 & 0 & 0 \\
0 & \nu & 0 & -1-\nu & 0 & 0 & 0 & 1-\nu & 0 & 0 & 0 & 1-\nu & 0 & 0 & 0 & 0 \\
\nu & 0 & 0 & 0 & -1 & 0 & 0 & 0 & 0 & 0 & 0 & 0 & 1-\nu & 0 & 0 & 0 \\
0 & \nu & 0 & 0 & 0 & -1 & 0 & 0 & 0 & 0 & 0 & 0 & 0 & 1-\nu & 0 & 0 \\
0 & 0 & \nu & 0 & 0 & 0 & -1 & 0 & 0 & 0 & 0 & 0 & 0 & 0 & 1-\nu & 0 \\
0 & 0 & 0 & \nu & 0 & 0 & 0 & -1 & 0 & 0 & 0 & 0 & 0 & 0 & 0 & 1-\nu \\
\nu & 0 & 0 & 0 & 0 & 0 & 0 & 0 & -(1-\nu) & 0 & 0 & 0 & 0 & 0 & 0 & 0 \\
0 & \nu & 0 & 0 & 0 & 0 & 0 & 0 & 0 & -(1-\nu) & 0 & 0 & 0 & 0 & 0 & 0 \\
0 & 0 & \nu & 0 & 0 & 0 & 0 & 0 & 0 & 0 & -(1-\nu) & 0 & 0 & 0 & 0 & 0 \\
0 & 0 & 0 & \nu & 0 & 0 & 0 & 0 & 0 & 0 & 0 & -(1-\nu) & 0 & 0 & 0 & 0 \\
0 & 0 & 0 & 0 & \nu & 0 & 0 & 0 & 0 & 0 & 0 & 0 & -(1-\nu) & 0 & 0 & 0 \\
0 & 0 & 0 & 0 & 0 & \nu & 0 & 0 & 0 & 0 & 0 & 0 & 0 & -(1-\nu) & 0 & 0 \\
0 & 0 & 0 & 0 & 0 & 0 & \nu & 0 & 0 & 0 & 0 & 0 & 0 & 0 & -(1-\nu) & 0 \\
0 & 0 & 0 & 0 & 0 & 0 & 0 & \nu & 0 & 0 & 0 & 0 & 0 & 0 & 0 & -(1-\nu)
\end{pmatrix}$$

#### B Timescales via an exponential growth model

To provide a timescale for the age of egg chambers, we fit an exponential growth model to measurements of the area of sections through egg chambers at different developmental stages. Area measurements used are from Table S2 in Shimada et al. [1].

#### C MCMC chains and convergence

Here we visualize in Figure S2 the MCMC chains obtained when sampling from the model at steady state and in the dynamic regime. Traceplots show the mixing of the chains and exploration of the space consistent across chains. Running mean plots indicate convergence of the mean value for each chain.

© 2019 The Authors

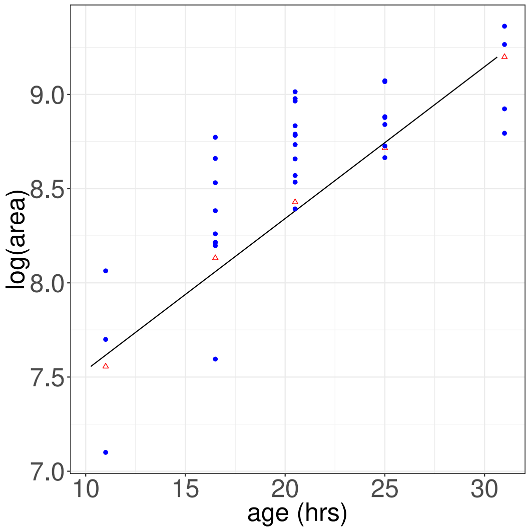


Figure S1: Fitting of exponential growth model to give timescale based on staging of egg chambers and measurements of area of the median section through the egg chamber. A linear model was used to fit the logarithm of the area as $\log(A) = \log(A_0) + \tau t$ via linear regression, as described in Section 2.5. The values of $\log(A_0)$ and $\tau$ found were 7.6 and 0.06, respectively. Red triangles show averaged data from Shimada et al. [1] used to fit the growth model, and blue points show measurements from egg chambers used.

### D Typical behaviour of the coarse-grained ODE model

Here we show typical behaviour from the coarse-grained ODE model across all the cells in the tissue in Figure S3.

### E Negative Binomial distribution parameterization

For the measurement model described in Section 3.2, we use a parameterization of the negative binomial distribution via a mean parameter, $\mu$, and overdispersion parameter, $\psi$, such that

$$p(y|\mu, \psi) = \binom{y + \psi - 1}{y} \left(\frac{\mu}{\mu + \psi}\right)^y \left(\frac{\psi}{\mu + \psi}\right)^\psi .$$

With this parameterization, if $y \sim \text{NB}(\mu, \psi)$, then $\mathbb{E}[Y] = \mu$ and $\text{Var}[Y] = \mu + \mu^2/\psi$.

### F Choice of prior distributions

The prior distributions are chosen to incorporate existing biological knowledge of feasible parameter ranges. For the rate parameters, $a$ and $b$, we choose Truncated Normal priors with parameters $\mu = 0$, $\sigma = 10$ and support $(0, \infty)$. Note that we define a Truncated Normal distribution with parameters $\mu$ and $\sigma$ and support on an interval $J \subset \mathbb{R}$ via the following rescaled probability density function

$$p(x; \mu, \sigma, J) = \frac{N(x; \mu, \sigma)}{1 - \int_{\mathbb{R} \setminus J} N(s; \mu, \sigma) \mathrm{d}s},$$

where $N(x; 0, 1)$ is the probability density function of a standard unit normal distribution. For the assembly parameter, $\phi$, we use a Truncated Normal prior with $\mu = 0.345$, $\sigma = 0.048$ and support $(0, 1]$. This is a strong prior informed by experimental evidence (see Section 3.3). For the transport bias parameter, $\nu$, we assume a uniform prior, $U(0, 0.5)$; it is clear that transport is biased towards the oocyte, but we avoid providing any further information about $\nu$. For the measurement noise parameter, $\sigma$,

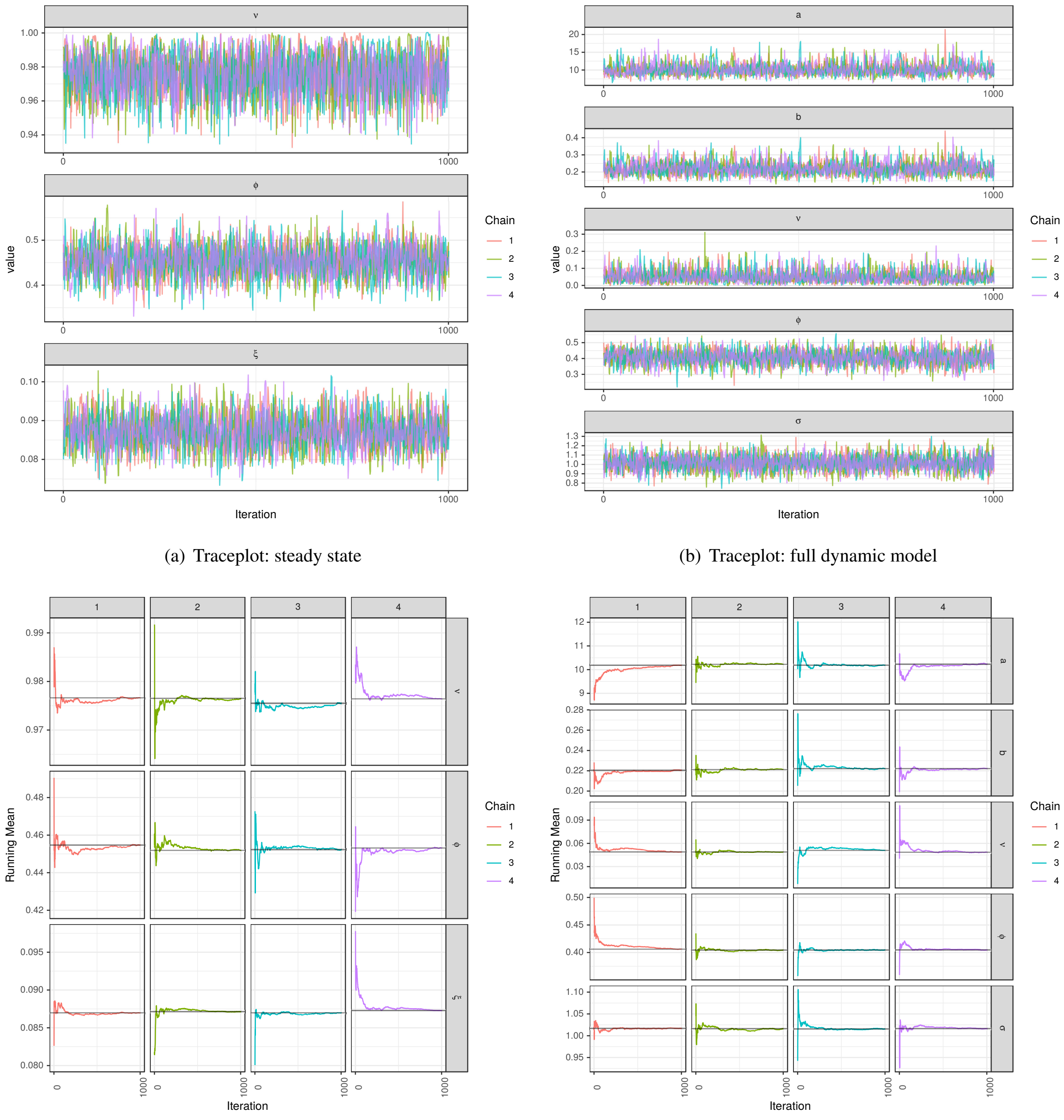


(a) Traceplot: steady state

(b) Traceplot: full dynamic model

(c) Running mean: steady state

(d) Running mean: full dynamic model

Figure S2: Convergence of MCMC chains for the model at steady state and the full model as described in Sections 3.4 and 3.5 respectively. Four Markov chains were run in parallel. Traceplots and running mean plots indicate convergence to the posterior distribution. Hamiltonian Monte Carlo implemented via Stan [2] was used to perform this sampling, as described in Section 2.6. The potential scale reduction statistic [3], $\hat{R}$, is 1.0 for inference at steady state, and also 1.0 for the full problem, indicating convergence in both cases. The minimum effective sample size, $n_{\text{eff}}$, across the model parameters is 1972 for inference at steady state, and 1914 for the full model.

we choose a broad Truncated Normal prior with $\mu = 0, \sigma = 10$ and support $(0, \infty)$. We check that these priors are compatible

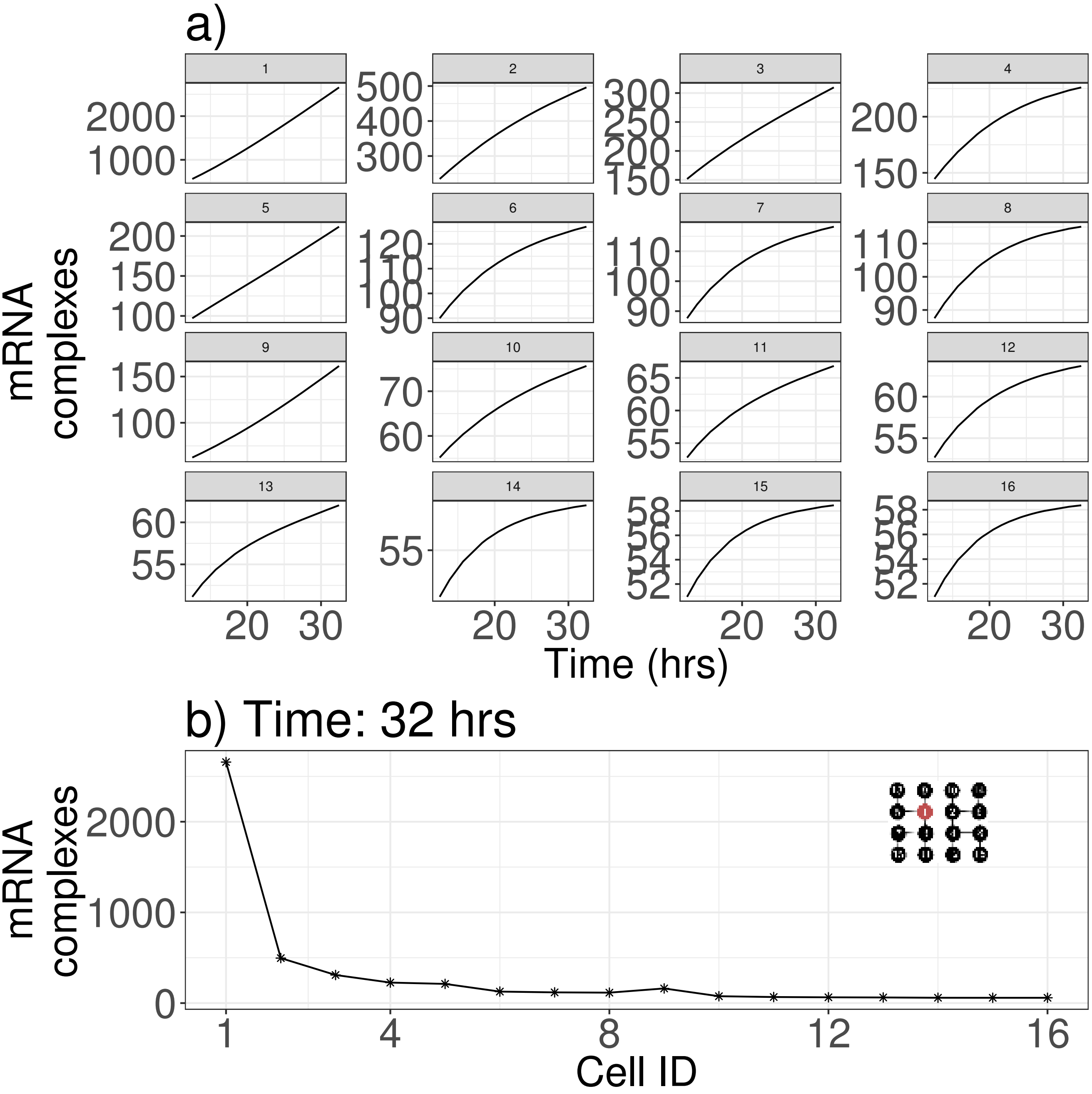


Figure S3: Typical behaviour of the coarse-grained ODE model across each of the cells in the egg chamber is shown in a). The distribution of mRNA across cells is shown in at a single time point at 32 hrs. The numbering for the Cell ID on the $x$ axis corresponds to that given in Figure 1. Parameters are $a = 10\,\text{particles hr}^{-1}$, $b = 0.20\,\text{hr}^{-1}$ and $\nu = 0.90$.

with the observed data by simulating from the prior predictive distribution, as suggested in the Bayesian workflow outlined by Gabry et al. [4] (see Figure S4).

For the model at steady state used in Section 3.4 to study the bias in transport through ring canals, we additionally have a prior on the measurement noise, $\xi$, in the form of a Truncated Normal prior with $\mu = 0$, $\sigma = 0.1$ and support $(0, \infty)$. Priors for $\nu$ and $\phi$ for the steady state model are as stated above.

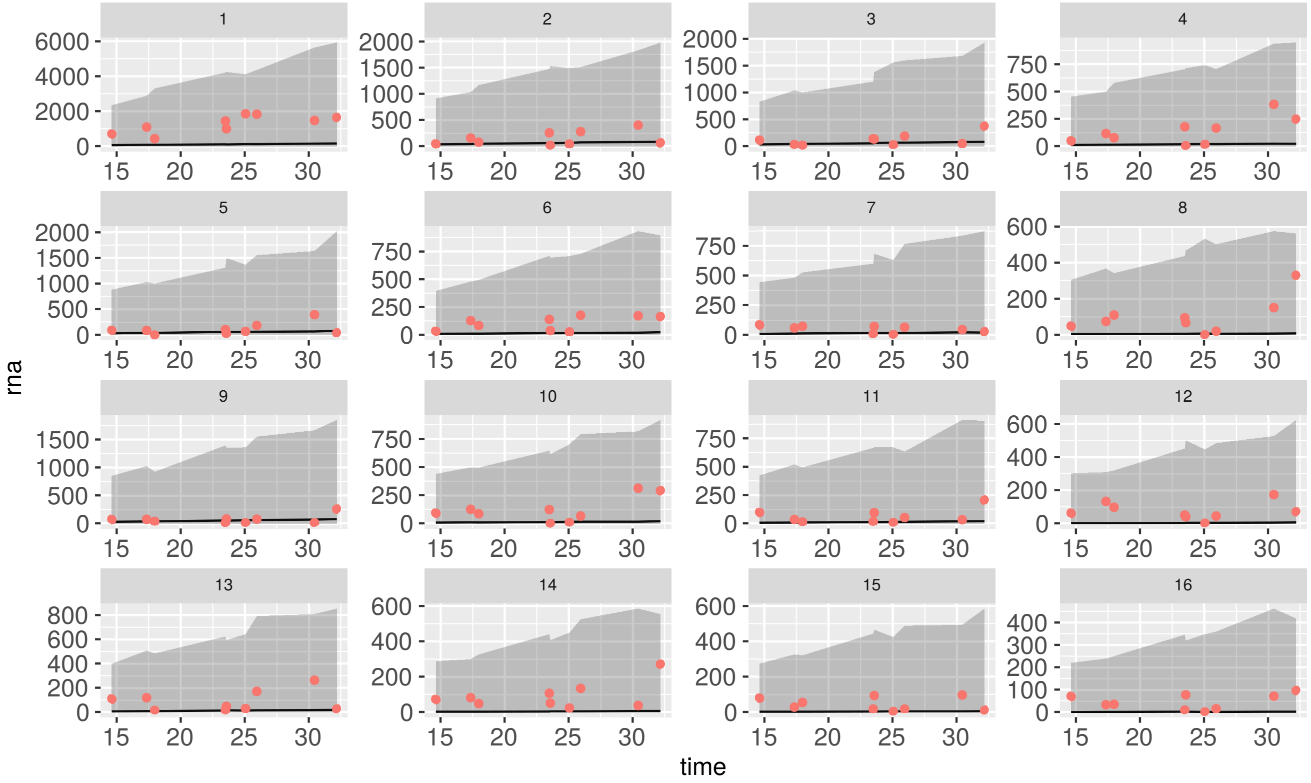


Figure S4: Predictions from the model given by Equation (1) before it has been fitted to the data, by drawing samples from the prior predictive distribution. Each separate facet plot gives predictions for a different nurse cell. The red dots show the observed wild type data. The shaded grey ribbon gives the 95% credible interval of predictions from the prior predictive distribution, while the black line gives the median prediction. The red dots clearly lie within the range of possible model predictions suggesting that the model and priors are flexible enough to generate the data.

### G Quantifying the mRNA content of individual RNA complexes

To investigate the mRNA content of individual RNA complexes and help quantify how many transcripts form a complex, we fitted a Gaussian mixture model to the background-subtracted intensity data for complexes in the nurse cells and oocyte. Following Little et al. [5], we normalized these intensities to the modal intensity value reasoning that this should represent a single transcript. The Gaussian mixture model is fitted via the Expectation-Maximization algorithm and reveals multimodal distributions of intensities with a two-component mixture model appropriate for the nurse cell data, while multiple components are needed for data from the oocyte. The mixture components lie at approximately integer values consistent with complexes formed from single or multiple transcripts.

### H Analytic solution to the coarse grained ODE model

The ordinary differential equation (ODE) model for the evolution of complex numbers in the egg chamber is given by

$$\frac{\mathrm{d}\mathbf{y}}{\mathrm{d}t} = a\mathbf{v} + b\mathbf{B}(\nu)\mathbf{y}, \tag{1}$$

where $\mathbf{y}$ gives the number of complexes in each of the cells of the egg chamber, $\mathbf{v}$ is a vector describing which cells produce mRNA, and $\mathbf{B}$ is a rate matrix where each row gives the relative rates at which RNA flows into and out of a cell. As an initial condition, we take $\mathbf{y}(t_0) = \mathbf{0}$.

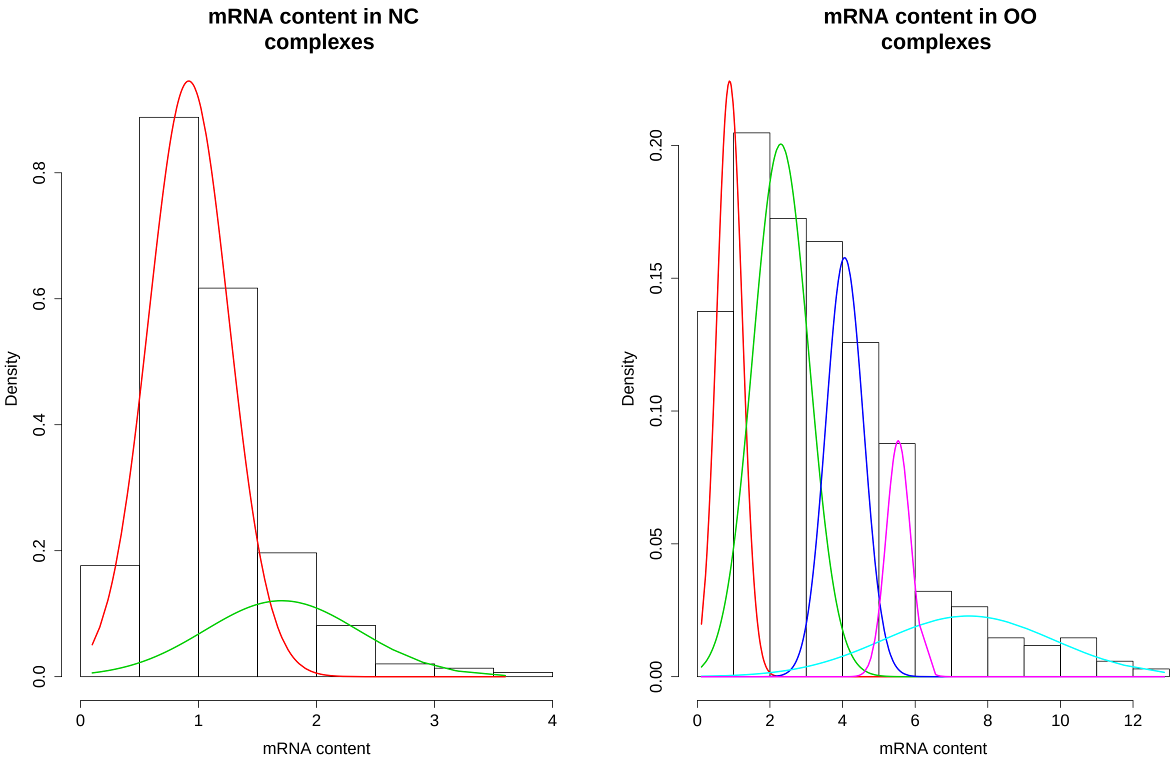


(a) (b)

Figure S5: Multimodal distribution of mRNA content of individual RNA complexes in the nurse cells and the oocyte. A Gaussian mixture model with components at integer values fits these data well, consistent with complexes formed from single and multiple transcripts.

Equivalently, we can write Equation (1) as

$$\mathrm{L}\mathbf{y} = f = a\mathbf{v},$$

where L is the linear operator

$$\mathrm{L}\mathbf{y} := \frac{\mathrm{d}\mathbf{y}}{\mathrm{d}t} - b\mathbf{B}(\nu)\mathbf{y}.$$

We note that we can solve the homogeneous problem $\mathrm{L}\mathbf{y} = 0$ by calculating the eigenvalues and eigenvectors of the matrix $b\mathbf{B}(\nu)$. The solution to the homogeneous problem is $\mathbf{y}_h = \mathbf{VDc}$, where $\mathbf{V}$ a matrix of eigenvectors of $b\mathbf{B}(\nu)$, $\mathbf{D}$ is a diagonal matrix with diagonal entries $e^{\lambda_i t}$, such that the $\lambda_i$ are the corresponding eigenvalues, and $\mathbf{c}$ is a vector of constants determined by the initial conditions.

For the particular solution, we take $\mathbf{y}_p = \mathbf{k_1} + \mathbf{k_2}t$, for some vectors $\mathbf{k_1}$, $\mathbf{k_2} \in \mathbb{R}^{16}$. Applying the linear operator to this particular solution gives

$$\mathrm{L}\mathbf{y}_p = \mathbf{k_2} - b\mathbf{B}\mathbf{k_1} - b\mathbf{B}\mathbf{k_2}t.$$

In addition, we note that the inhomogeneous problem, $\mathrm{L}\mathbf{y} = a\mathbf{v}$, has a solution that has a constant term, but no term linear in $t$. Therefore, we must have $\mathbf{k_2} \in \ker(\mathbf{B})$. Provided that $\nu \in [0, 1]$, this allows us to find $\mathbf{k_2}$ uniquely (up to a constant of proportionality), since the null space of $\mathbf{B}$ has dimension 1. That the null space has dimension 1 is due to the zero eigenvector of $\mathbf{B}$, which is present due to that fact that no complexes are lost from the system (we can also use this fact to determine the constant of proportionality).

Given we can determine $\mathbf{k_2}$ by finding the null space of B, we can solve the following for $\mathbf{k_1}$:

$$-b\mathbf{B}\mathbf{k_1} = a\mathbf{v} - \mathbf{k_2}. \tag{2}$$

Since $\mathbf{B}$ is not a full rank matrix, we cannot uniquely determine a solution to Equation (2); instead we obtain a line of solutions of the form $\mathbf{k_1} = \mathbf{k_1}^* + \lambda \mathbf{k_2}$, for some parameter $\lambda$.

The constants in the vector $\mathbf{c}$ can be determined from the initial conditions, $\mathbf{y_0}$. Initially at $t = 0$,

$$\begin{aligned} \mathbf{y_0} &= \mathbf{V}\mathbf{c} + \mathbf{k_1}, \\ \implies \mathbf{c} &= \mathbf{V}^{\mathsf{T}}(\mathbf{y_0} - \mathbf{k_1}^* - \lambda\mathbf{k_2}). \end{aligned}$$

Put together, we have a general solution for the ODE model of the form

$$\mathbf{y} = \mathbf{V}\mathbf{D}\mathbf{c} + \mathbf{k_1} + 15a\mathbf{k_2}t. \tag{3}$$

### I Pairwise posterior distribution

After fitting the full dynamic ODE model (3.2), we obtain a posterior distribution across all the model parameters. To visualize this distribution, we show pairwise and marginal plots for all the model parameters in Figure S6.

### J Sensitivity to rate parameters

We provide further evidence to support the hypothesis that production and transport are in tightly regulated balance by performing a sensitivity analysis of the model described in Section H. We consider the effects of making a small perturbation to the production and transport rates $a$ and $b$. Since Equation (1) is a linear system of ODEs, we can obtain an analytical solution for the evolution of $\mathbf{y}$, as given in Equation (3). By differentiating this solution with respect to model parameters $a$ and $b$, we find an expression for how sensitive the number of RNA-protein complexes is to these parameters. In the long time limit, $\partial\mathbf{y}/\partial a$ is linear in time, whereas $\partial\mathbf{y}/\partial b$ tends to a constant. Since biological transport behaviour occurs in this long time limit, we have that dependence of the number of complexes on production rate, $a$, dominates the dependence on transport rate, $b$.

Using the general solution obtained in Equation (3), and differentiating with respect to the production rate, $a$, we have

$$\begin{aligned} \frac{\partial\mathbf{y}}{\partial a} &= \mathbf{V}\mathbf{D}\frac{\partial\mathbf{c}}{\partial a} + \frac{\partial\mathbf{k_1}}{\partial a} + \frac{\partial\mathbf{k_2}}{\partial a}t, \\ &= (I - \mathbf{V}\mathbf{D}\mathbf{V}^{-1})\frac{\partial\mathbf{k_1}}{\partial a} + \frac{\partial\mathbf{k_2}}{\partial a}t. \end{aligned}$$

We obtain a similar expression when differentiating with respect to the transport rate, $b$, but without the term giving linear growth in time

$$\frac{\partial\mathbf{y}}{\partial b} = (I - \mathbf{V}\mathbf{D}\mathbf{V}^{-1})\frac{\partial\mathbf{k_1}}{\partial b}.$$

We show in Figure S7 how sensitivity to the rate of production, $a$, and the rate of transport, $b$, vary across different parameter values, and highlight relevant biological parameters by the orange circle. At realistic parameter values: $(a, b) = (10, 0.2)$, then

$$\frac{\partial\mathbf{y}}{\partial a} \approx \frac{\partial\mathbf{y}}{\partial b}.$$

### K Model comparison

Models can be evaluated based on the quality of their predictions about future behaviour, based on unknown future observations, $\tilde{\mathbf{y}}$ [6]. The posterior predictive distribution for a model, $M$, based on observed data, $\mathbf{y}$, can be written as:

$$p(\tilde{\mathbf{y}}|\mathbf{y}, M) = \int p(\tilde{\mathbf{y}}|\theta, M)p(\theta|\mathbf{y}, M)\,\mathrm{d}\theta, \tag{4}$$

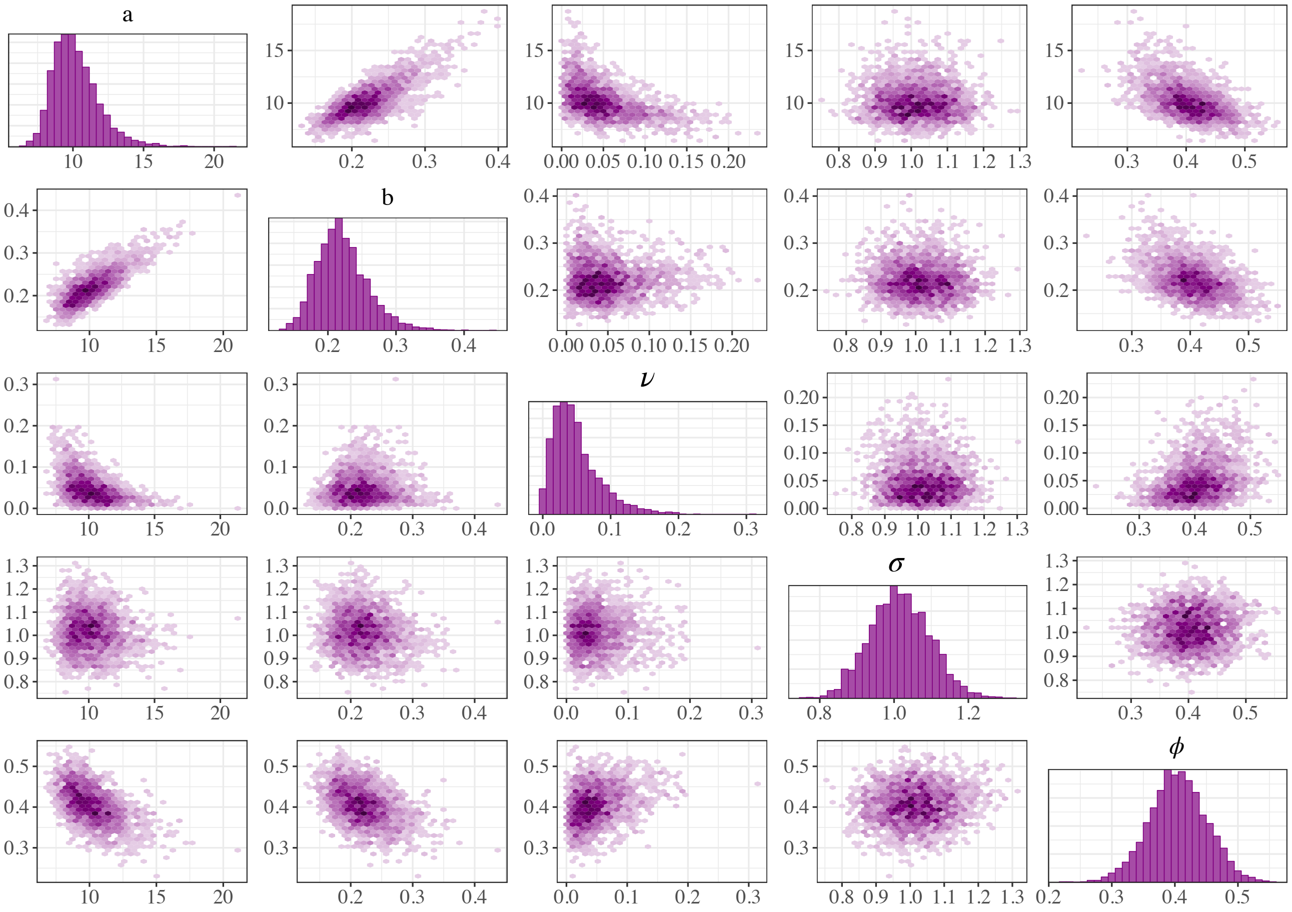


Figure S6: Pairwise posterior plot for the full dynamic coarse-grained ODE model (3.2). Marginal distributions for each parameter ($a$, $b$, $\nu$, $\sigma$, $\phi$) are shown on the diagonal. Pairwise plots are shown as hex plots with darker purple indicating higher density. We note the positive correlation between production rate $a$ and transport rate $b$ suggesting that these rates are in a tightly regulated balance.

by marginalizing out the model parameters, $\theta$. Evaluation of the quality of predictions requires a utility function, $u(M, \tilde{\mathbf{y}})$ [7]. A common choice for this utility function is the logarithmic score

$$u(M, \tilde{\mathbf{y}}) = \log p(\tilde{\mathbf{y}}|\mathbf{y}, M), \tag{5}$$

which possesses desirable properties, such as invariance to reparameterization [8], and a connection to the Kullback-Leibler (KL) divergence between the true distribution of $\tilde{\mathbf{y}}$ and the distribution based on predictions from model $M$ [7].

Consideration of the log score as a utility function leads to the following Monte Carlo estimator for the expected log predictive distribution (ELPD) [9]:

$$\mathbb{E}\left[\log\left(\int p(\tilde{\mathbf{y}}|\theta)p(\theta|\mathbf{y})\,\mathrm{d}\theta\right)\right] \approx \frac{1}{n}\sum_{i=1}^{n}\log\left(\int p(y_i|\theta)p(\theta|\mathbf{y})\,\mathrm{d}\theta\right). \tag{6}$$

Cross validation allows the predictive performance of a model to be assessed by efficient use of a single dataset, rather than requiring collection of a full new independent dataset on which to assess predictive performance. A dataset is divided

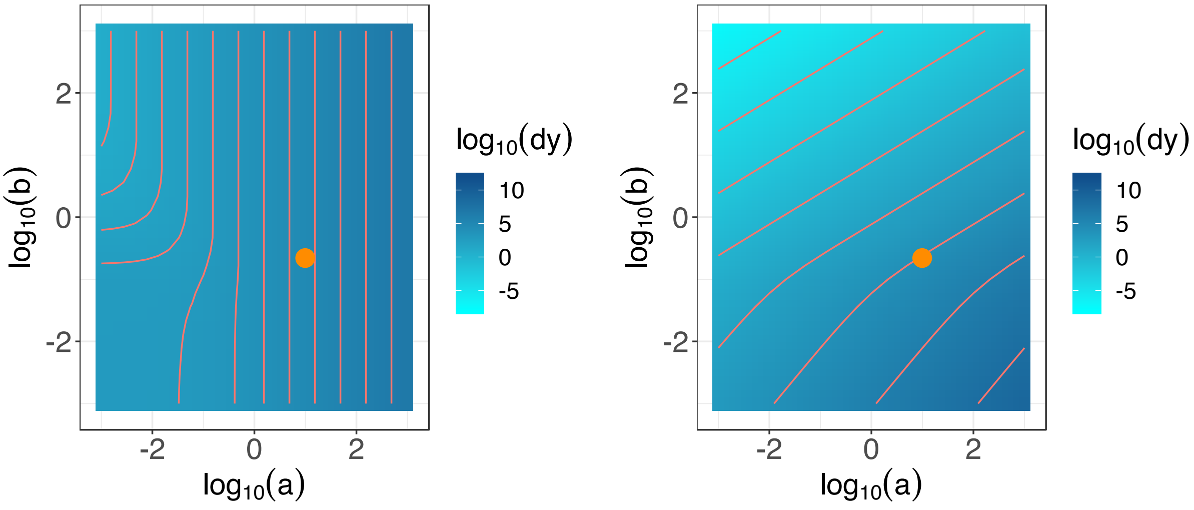


(a) Sensitivity of $\frac{\partial y_1}{\partial a}$ (b) Sensitivity of $\frac{\partial y_1}{\partial b}$

Figure S7: Sensitivity analysis for the ODE model given in (1). In (a), we show the sensitivity of the number of RNA particles in the oocyte to the rate of production, $a$, with realistic model parameters highlighted with an orange circle. Similarly in (b), we show sensitivity to the rate of transport, $b$. Results are presented at time $t = 30$ hrs.

into equally sized subsets, one of which is retained to evaluate the performance of a model fitted to the remaining data. For example, from a dataset of $n$ points of observed data $\mathbf{y}$, we can produce $n$ subsets of the data $\mathbf{y}_{-i}$ for which the single point $y_i$ has been removed. Using these subsets to evaluate predictive performance of a model results in leave-one-out cross validation [10, 11].

Weights for a collection of models under comparison can be obtained via the ELPD defined in Equation (6). Using a Bayesian leave-one-out estimator of the ELPD [12] gives

$$\mathrm{ELPD}_{\mathrm{loo}} = \sum_{i=1}^{n} \log(p(y_i|\mathbf{y}_{-i})) \tag{7}$$

where $p(y_i|\mathbf{y}_{-i}) = \int p(y_i|\theta)p(\theta|\mathbf{y}_{-i})\, \mathrm{d}\theta$ is the leave-one-out predictive density omitting the $i$th data point. Suppose $\mathrm{ELPD}^j_{\mathrm{loo}}$ is the estimated expected log predictive density for the model $M_j$, then the corresponding weight $w_j$ for model $M_j$ can be computed as

$$w_j = \frac{\exp\left(\mathrm{ELPD}^j_{\mathrm{loo}}\right)}{\sum_{j=1}^{J} \exp\left(\mathrm{ELPD}^j_{\mathrm{loo}}\right)}. \tag{8}$$

These weights are known as Pseudo-BMA (Bayesian model averaging) weights. The Pseudo-BMA weights can provide an assessment of which model is most predictive from a set of models. Bayesian bootstrap methods can compute uncertainties for the leave-one-out estimates based on the $n$ independent components from different data points [9], which leads to adjusted weights known as Pseudo-BMA+ weights.

#### *K.1 Stacking*

Recent work has proposed using stacking of predictive distributions to combine predictive distributions from different models together and aid in model comparison [9]. This approach is based on previous work on averaging of point estimates for multiple models [13, 14]. This technique can help avoid some of the criticisms of Bayesian model averaging and other model comparison methods.

In the stacking method of Yao et al. [9], the log score of the combined predictive distribution is maximized subject to the weights combining predictive distributions for each model being positive, normalized weights:

$$\max_{\mathbf{w}} \frac{1}{n}\sum_{i=1}^{n} \log\left(\sum_{j=1}^{J} w_j p(y_i|y, M_j)\right) \quad \text{subject to } w_j \geq 0, \sum_{j=1}^{J} w_j = 1, \tag{9}$$

where $w_k$ are the stacking weights for each model. Asymptotically, rather than concentrating on a single model closest to the true model in KL divergence, the stacking weights method will find the optimal predictive distribution within the convex set formed from the span of models. This may be more informative in the case where the true model is not contained in the set of models considered.

### L Estimates of $\gamma$

We assume that the over-expression mutant has RNA complex production rate $\gamma a$, where $a$ is the production rate in wild type, and $\gamma > 1$ is a scale factor. We estimate $\gamma$ by considering the total mRNA over time in all cells in the egg chamber (15 nurse cells and the oocyte). Recall that mRNA is produced in the nurse cells, but not the oocyte which is transcriptionally silent, such that if we define the total mRNA across all cells as:

$$Y(t) = \sum_{i=1}^{16} y_i(t), \tag{10}$$

then by summing Equation (1), the transport terms cancel and we obtain

$$\frac{\mathrm{d}Y}{\mathrm{d}t} = 15a. \tag{11}$$

Suppose that we observe $Z_{\mathrm{WT}} \sim \mathcal{N}(Y_{\mathrm{WT}}, \sigma^2_{\mathrm{WT}})$ and $Z_{\mathrm{OE}} \sim \mathcal{N}(Y_{\mathrm{OE}}, \sigma^2_{\mathrm{OE}})$ for the wild type and over-expression mutant, respectively, where we assume that $Z_{WT}$ and $Z_{OE}$ are the total observed mRNA counts, and assume production at rates $a_{\mathrm{WT}}$ and $a_{\mathrm{OE}}$, with $a_{\mathrm{OE}} = \gamma a_{\mathrm{WT}}$.

We assume the following prior distributions: Truncated Normal distribution with $\mu = 0$, $\sigma = 10$ and support $(0, \infty)$ for the production rates $a_{\mathrm{OE}}$ and $a_{\mathrm{WT}}$; Half-Cauchy$(0, 3)$ with support on $(0, \infty)$ for observation noise parameters $\sigma_{\mathrm{OE}}$ and $\sigma_{\mathrm{WT}}$. Results of performing MCMC for this model are shown in Figure S8, giving a median estimate for $\gamma$ of 2.23, with a 95% credible interval of $[1.38, 3.06]$.

### M Dependence of distribution of mRNA on distance from the oocyte

Previous work [15] has shown that intercellular connections are sufficient to explain the correlation of nurse cell volumes with distance from the oocyte. Here we illustrate the dependence of the distribution of mRNA in each nurse cell on its distance from the oocyte within the graph structure of Figure 1c). Dependence on the distance from the oocyte is not immediately clear from the noisy raw mRNA counts, but some structure and groupings of cells can be seen when the mRNA counts are ranked from 1 to 16 for each egg chamber, where 1 corresponds to the highest mRNA count. In particular, the ranking increases with distance from the oocyte in wild type. This structure is altered in the over-expression phenotype due to spikes of higher mRNA counts in some cells far from the oocyte resulting in lower ranks.

### N Quantifying production via nascent transcription

Previously in Section 3.1, we assumed equal production of mRNA in each of the nurse cell nuclei. Here we relax this assumption and consider estimating directly how much production occurs in each nurse cell nucleus. The UAS system is used as a

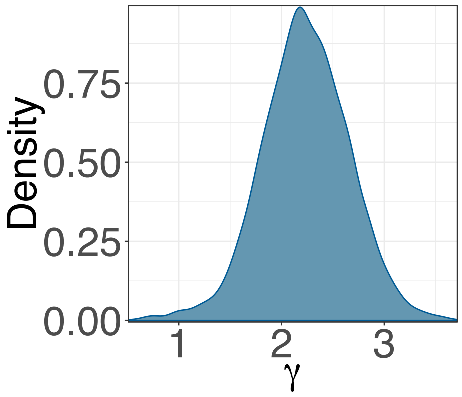


Figure S8: Posterior distribution for the ratio, $\gamma$, between production in wild type and the over-expression mutant.

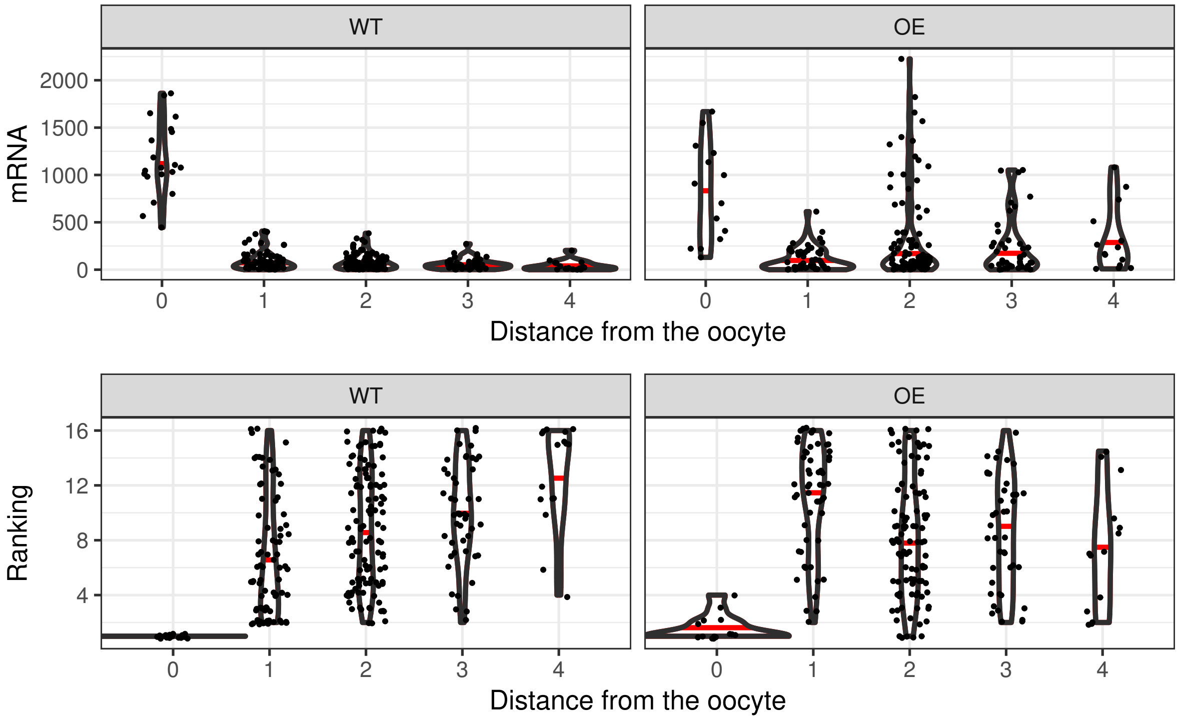


Figure S9: Dependence of distribution of mRNA on distance from the oocyte in both wild type and the over-expression mutant. Differences between nurse cells at different distances from the oocyte are not clear based on the raw mRNA counts, but when the mRNA counts for cells within an egg chamber are ranked from 1 to 16, structure is revealed in the data reflecting the graph of connections between cells. The ranking increases with distance from the oocyte in wild type. However, due to much higher mRNA counts in some nurse cells in the over-expression mutant, this structure is obscured in the over-expression mutant. Violin plots show the distribution of data marked by the dots, with the horizontal red bar marking the median of the distribution.

genetic driver to drive the over-expression of *gurken*, and this may not operate identically in all the nurse cells. This justifies considering inhomogeneous production for the over-expression mutant, but nonetheless assuming equal production across nurse cells in the wild type.

Challenges arise in the nature of the data available in that the smFISH data is from fixed specimens and so captures a snapshot of the dynamics at a particular point in time. Transcription is a noisy process and there is some evidence that time averaging of transcription helps regulate gene expression [16]. By considering only a single sample in time from each nurse cell, we lack information to make sound inferences that can extrapolate to other time points. Instead, we assume that there is a characteristic distribution of transcription across cells within the egg chamber.

To quantify transcription in each nurse cell, we measure the brightness of nascent transcripts in the nuclei of nurse cells. We perform these measurements based on an average projection in the $z$ direction through the nucleus of each nurse cell, and apply a threshold to remove background signal using FIJI (ImageJ V1.51d; $fiji.sc$, [17]). The total intensity measurements are then scaled by the total area of foci in the nucleus. We note that the resulting distributions of fluorescence across cells for different over-expression egg chambers are very noisy with large spikes, as shown in Figure S10, whereas in wild type, although the distributions are still noisy, they are closer to constant production across nurse cells.

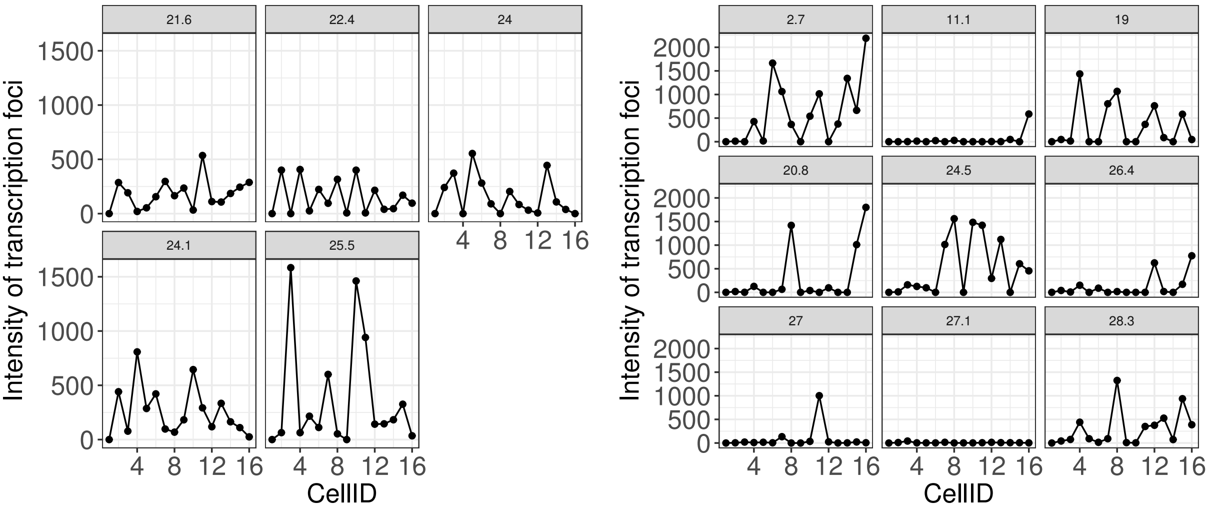


(a) Nascent transcription in WT

(b) Nascent transcription in OE

Figure S10: Nascent transcription measured in transcription foci in the nuclei of nurse cells from *Drosophila* egg chambers in wild type ((a)), and with a *gurken* over-expression phenotype ((b)). Each subplot corresponds to a different egg chamber, with the title indicating the age of the egg chamber in hours. The index for each cell shown on the $x$ axis corresponds to the numbering of the graph in Figure 1. The transcription shown on the $y$ axis is in units of fluorescence.

We note that the measured nascent transcription rates in wild type are consistent with the simplest modelling assumption of equal production across nurse cells. This assumption provides additional benefits of tractability and simplicity. Although we cannot exclude the possibility that a different inhomogeneous distribution of production applies here, there is not sufficient evidence for it based on the available snapshots at single times. For these reasons we have used the assumption of equal production in wild type.

For the over-expression mutant, in order to consider an inhomogeneous production model, we infer a relative production rate for each cell. A notable minority of nurse cells for the over-expression mutant have a measured nascent transcription value of zero after removal of background. To overcome problems caused from these zero observations, we use a mixture model and treat the data as missing with some probability, $p$, in which case we observe zero for the nascent transcription in a given cell; otherwise with probability $1-p$ we use a poisson model for the nascent transcription. Suppose $\mathbf{a} = (a_2, \ldots, a_{16})$

are the production rates in each cell (we assume $a_1 = 0$ since the oocyte is transcriptionally silent), and suppose also that $\tau = (\tau_2, \ldots, \tau_{16})$ are the observed nascent transcription values. The model to estimate the rates of inhomogeneous production in each cell takes the following form

$$\tau \sim (1-p)\,\text{Poisson}(\tilde{\mathbf{a}}) + p\,\delta(x), \tag{12}$$

where $\delta(x)$ is a delta function, and $\tilde{\mathbf{a}}$ are the unnormalized production rates such that $\mathbf{a} = 15\gamma\tilde{\mathbf{a}}/\sum_i \tilde{a}_i$. We assume a prior uniform on $(0, 1)$ for the missing data probability, $p$, and a Half-Cauchy$(0, 5)$ prior with support on $(0, \infty)$ for the inhomogeneous production rate $\tilde{\mathbf{a}}$. The resulting estimates for inhomogeneous production, $\mathbf{a}$, across nurse cells of the egg chamber are shown in Figure S11 and are used in Section Q for the inhomogeneous production model.

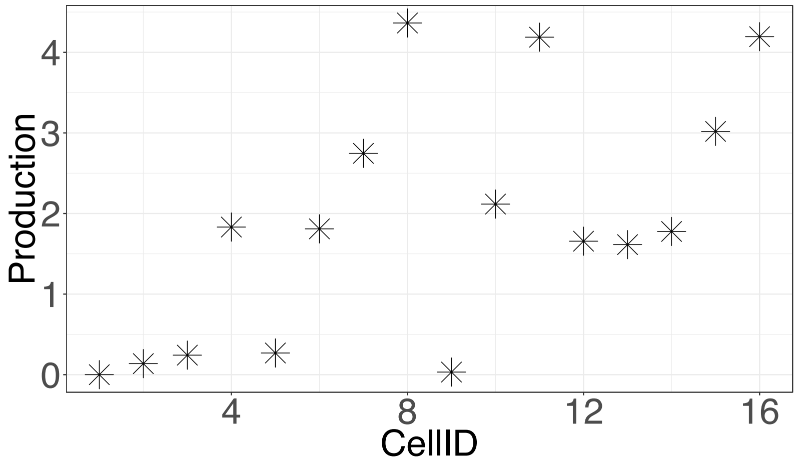


Figure S11: Estimated inhomogeneous production values across cells based on measurements of nascent transcription and a mixture model. Points shown are median estimates for the production in each cell. Posterior distributions are tight around these median values. Note that the normalized production values are shown such that an homogeneous model for the over-expressor would have production of $\gamma$ across all nurse cells, and the wild type would have production of 1 in each nurse cell.

### O Quantification of blocking of ring canals

We show the distribution of mRNA counts for each egg chamber both in wild type and for the over-expression mutant in Figure S12. These profiles of mRNA counts across cells can indicate build up of mRNA in certain cells linked to blocking behaviour. Additionally, the mRNA count profiles across cells highlight differences between the wild type and over-expression phenotype.

### P Extension to model with mRNA decay

We consider a generalization of the model described in Equation (1) and Section 3.2 that accounts for degradation of mRNA. We assume that mRNA degrades at rate $\delta$ in each of the cells. Including decay in the model fundamentally changes the dynamics of the system of equations. In the quasi-steady-state, the mRNA levels in the nurse cells will be constant when decay is included whereas without decay mRNA levels in the nurse cells increase linearly over time.

The resulting system of equations with degradation of mRNA at rate $\delta$ is as follows:

$$\frac{\mathrm{d}\mathbf{y}}{\mathrm{d}t} = a\,\mathbf{v} + b\,\mathbf{B}(\nu)\,\mathbf{y} - \delta\,\mathbf{y}. \tag{13}$$

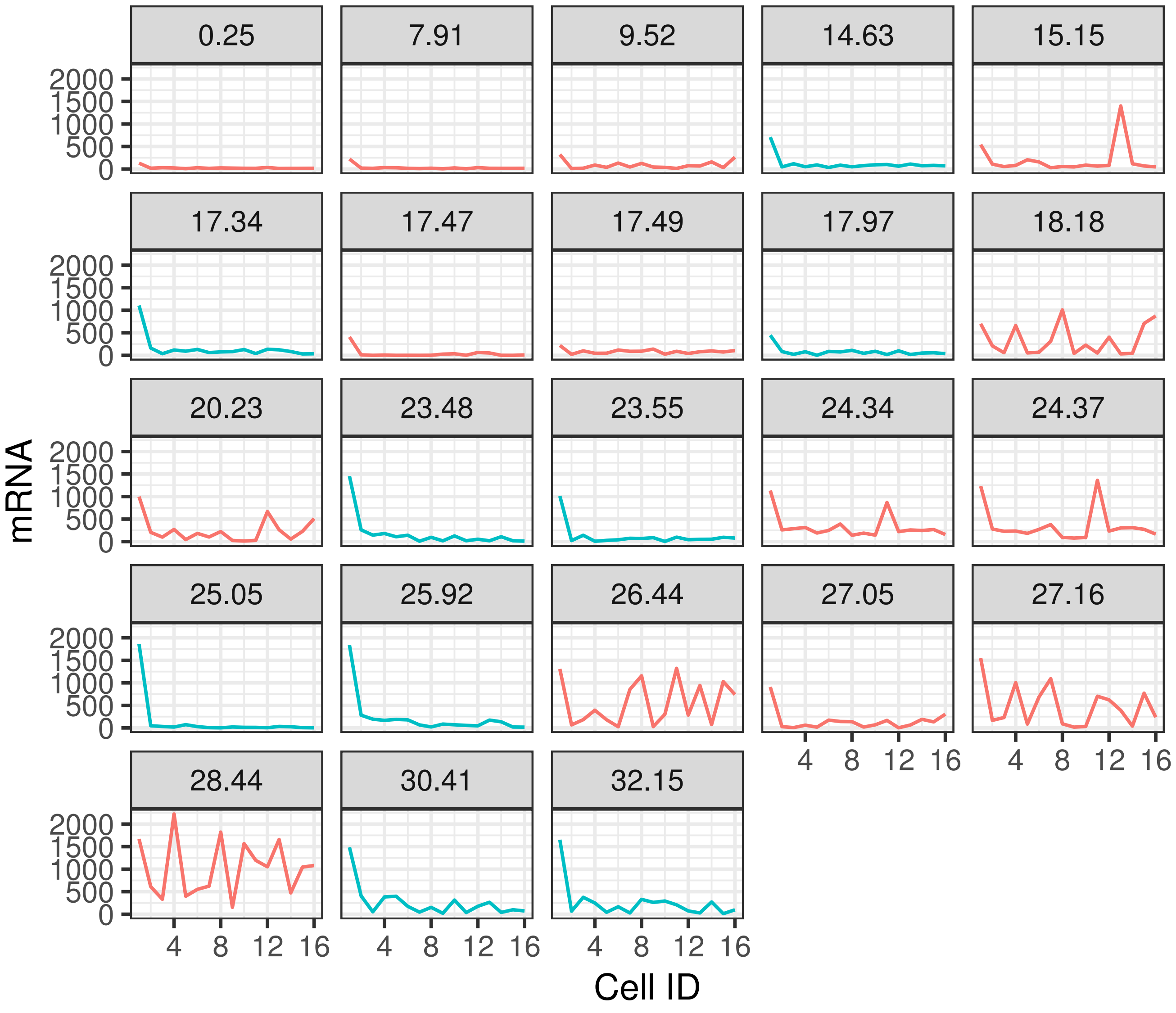


Figure S12: Distribution of mRNA counts across cells for each egg chamber. Data from the over-expression mutant is shown in red and from wild type in blue. Spikes in mRNA counts in cells far from the oocyte suggest blocking behaviour in these cells. Each subplot shows data from a different egg chamber with the title of each subplot indicating the age in hours based on the exponential growth model.

Setting the same prior for the decay rate, $\delta$, as for the other rate parameters of $\mathcal{N}(0, 10^2)$ in units of $\left[\text{hr}^{-1}\right]$, and otherwise using the same priors described in Section F, we can perform inference for the model with degradation given by Equation (13). We obtain a 95% credible interval for $\delta$ of [0.01,0.33] with a median of 0.14. Samples from the posterior predictive distribution for this model are shown in Figure S13. Conclusions drawn from our analyses are unaffected by using the model with decay.

An alternative hypothesis to explain the lack of agreement between predictions from the simple model and data from the over-expression phenotype is that degradation of RNA in the over-expression mutant occurs at a different rate to degradation

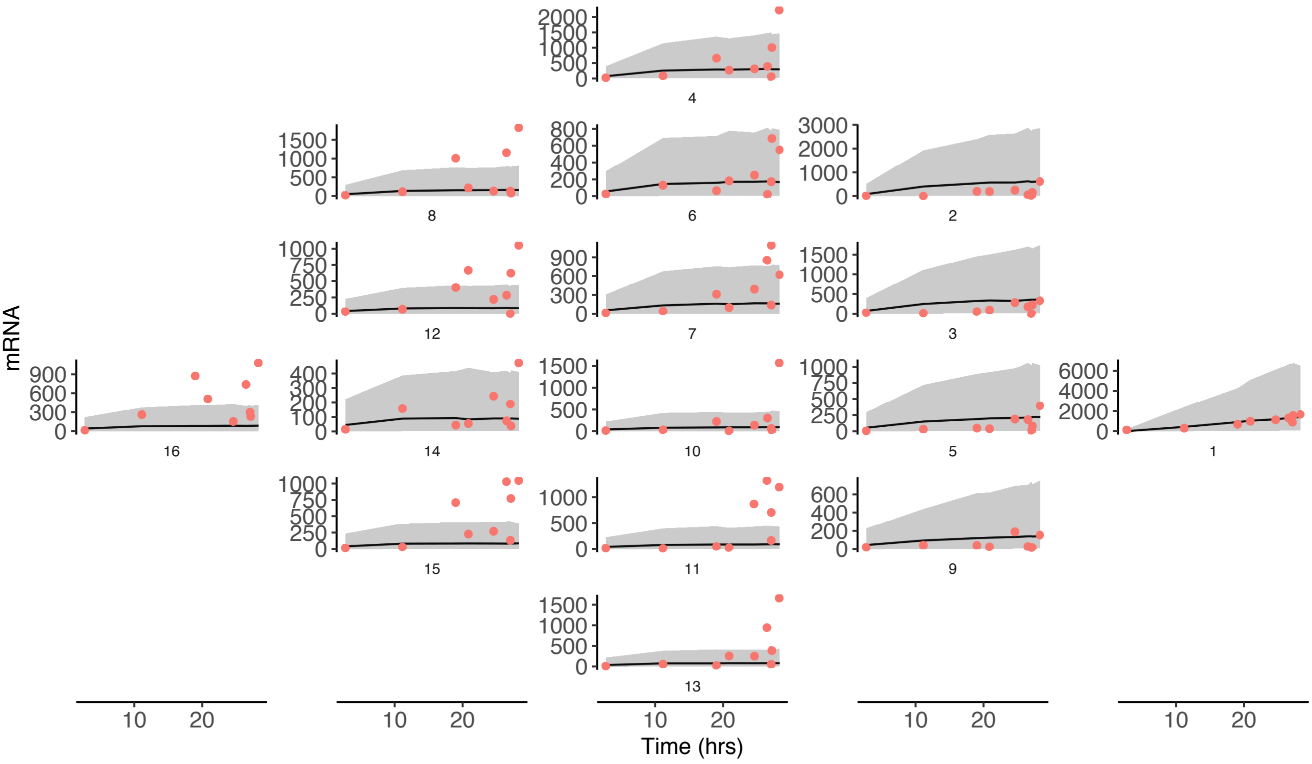


Figure S13: Posterior predictive distribution for the over-expression mutant using the model with mRNA decay at rate $\delta$. The gray envelope shows a 95% credible interval for predictions, while the red points correspond to observed data.

of RNA in wild type. In Section 3.8, we assessed competing models by fitting them to data from wild type and making a prediction about behaviour for the over-expression phenotype. For this hypothesis regarding the degradation of RNA, we require data from which to determine the degradation rate for the over-expression phenotype based only on wild type data, which is not available. We must instead determine this rate by fitting to over-expression data. Although doing so offers a favourable fit to the observed over-expression data, it does not provide a fair comparison to the other models since we would be learning from the over-expression data and assessing our predictions on it also ($p(\mathbf{y}_{OE}|\mathbf{y}_{WT}, \mathbf{y}_{OE})$ versus $p(\mathbf{y}_{OE}|\mathbf{y}_{WT})$). However, we note that degradation of RNA is an important regulatory mechanism in general, and we cannot rule out a role for it here.

### Q Analysis of inhomogeneous variance model

Previously we have assumed a homogeneous noise model across cells in the egg chamber, in that the overdispersion parameter, $\sigma$, in the negative binomial measurement model in Equation 2 is the same for every cell. The observed data, shown as dots in Figures 5 and 6, seems to exhibit greater variation in certain cells than others. We attempt to account for this variation by relaxing the assumption that the overdispersion parameter is the same for all cells and instead letting it vary across the tissue. Predictions for the over-expression mutant from the simple model, M0, but with inhomogeneous overdispersion, $\sigma$, are shown in Figure S14. The biggest difference is a smaller range of predictions for the oocyte. This is consistent with the robustness of the amount of mRNA localized in the oocyte. The disparity between the prediction for the over-expression mutant and the observed levels of mRNA localization in the oocyte are clearly apparent from these results also.

Further to this we considered a comparison of the collection of models $\mathcal{M} = \{\mathrm{M0}, \mathrm{M1}, \ldots, \mathrm{M7}\}$ between the homogeneous and inhomogeneous overdispersion cases. We computed weights jointly for this collection of 16 models together. The

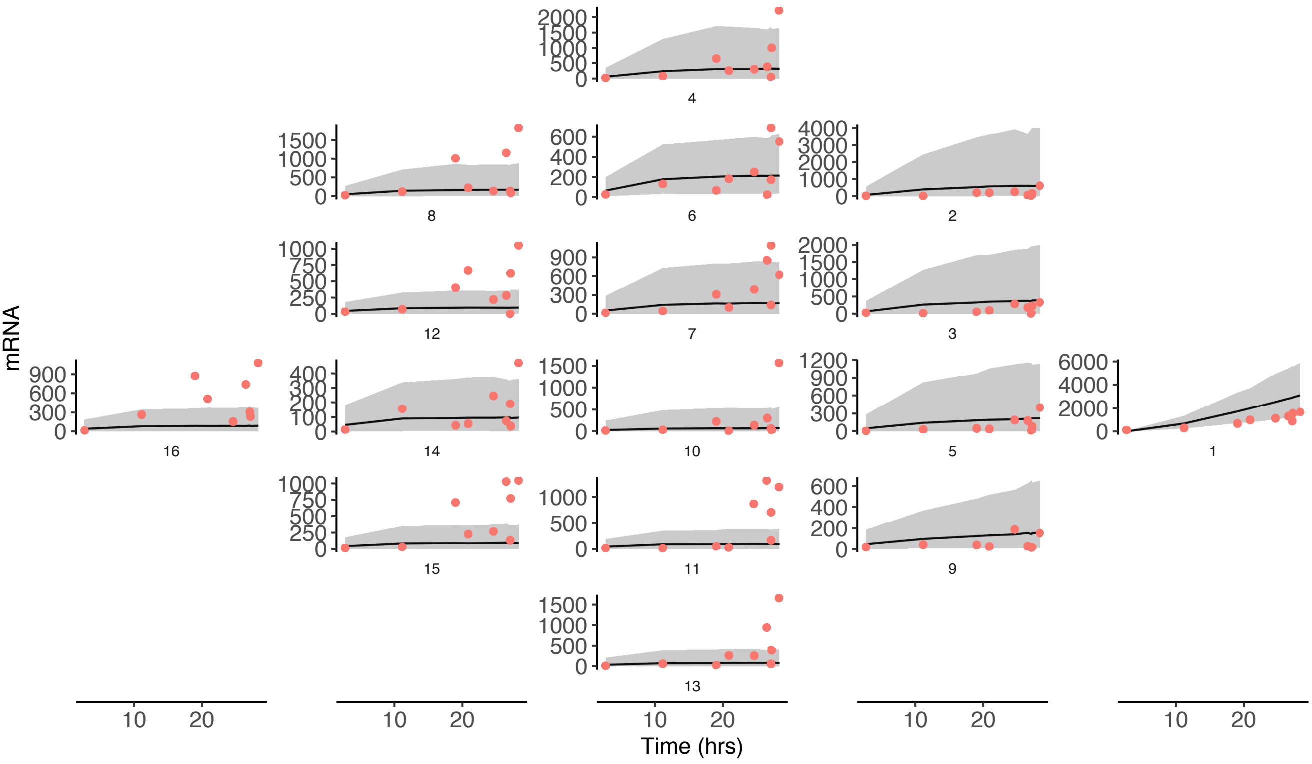


Figure S14: Posterior predictive distribution for the over-expression mutant using the overdispersion model, where a separate overdispersion parameter has been fitted in the measurement model for each cell in the egg chamber. The gray envelope shows a 95% credible interval for predictions, while the red points correspond to observed data.

results are shown in Figure S15 and demonstrate that, in general, the homogeneous overdispersion parameter models should be favoured over the inhomogeneous overdispersion parameter models. This justifies using the model with homogeneous overdispersion across cells. Moreover, the conclusion that crowding-induced blocking of ring canals is the most plausible model to explain the observed data for the over-expression mutant still holds when jointly comparing this larger collection of models. We note that the inhomogeneous overdispersion model is a more flexible class of model, but that this overfits to the wild type data and is less able to make predictions relevant for the over-expression mutant. This could indicate further biological differences between the wild type and over-expression mutant, which could be investigated in future studies.

### References

[1] Shimada, Y., K. M. Burn, R. Niwa, and L. Cooley, 2011. Reversible response of protein localization and microtubule organization to nutrient stress during Drosophila early oogenesis. Developmental Biology 355:250–262.

[2] Carpenter, B., A. Gelman, M. Hoffman, D. Lee, B. Goodrich, M. Betancourt, M. Brubaker, J. Guo, P. Li, and A. Riddell, 2017. Stan: A Probabilistic Programming Language. Journal of Statistical Software 76:1–32.

[3] Gelman, A., and D. B. Rubin, 1992. Inference from iterative simulation using multiple sequences. Statistical Science 7:457–472.

[4] Gabry, J., D. Simpson, A. Vehtari, M. Betancourt, and A. Gelman, 2017. Visualization in Bayesian workflow. arXiv preprint arXiv:1709.01449 .

[5] Little, S. C., K. S. Sinsimer, J. J. Lee, E. F. Wieschaus, and E. R. Gavis, 2015. Independent and coordinate trafficking of single Drosophila germ plasm mRNAs. Nature Cell Biology 17:558.

[6] Vehtari, A., and J. Ojanen, 2012. A survey of Bayesian predictive methods for model assessment, selection and comparison. Statistics

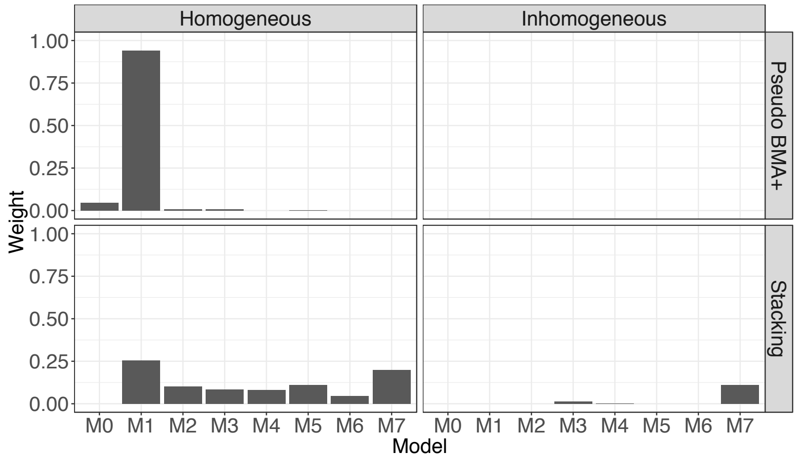


Figure S15: Model comparison weights between homogeneous and inhomogeneous overdispersion models based on a joint comparison of 16 models. Weights are computed via both the pseudo-BMA+ and stacking methods.

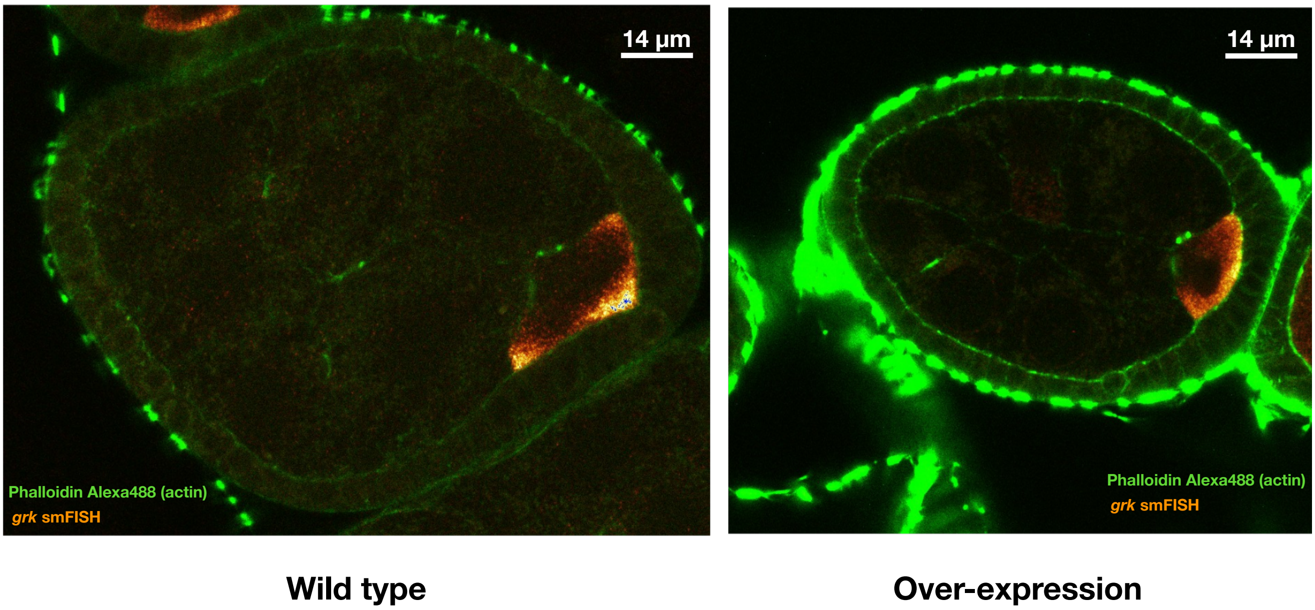


Figure S16: Comparison of phenotypes between wild type and over-expression mutant for egg chambers at similar stages of development and under identical imaging conditions. Similar amounts of *gurken* mRNA have been localized in each case. For the over-expression mutant, one of the nurse cells contains much more mRNA than the others, whereas the distribution across nurse cells for the wild type is more equal.

Surveys 6:142–228.

[7] Piironen, J., and A. Vehtari, 2017. Comparison of Bayesian predictive methods for model selection. Statistics and Computing 27:711–735.

[8] Robert, C. P., 1996. Intrinsic losses. Theory and Decision 40:191–214.

[9] Yao, Y., A. Vehtari, D. Simpson, and A. Gelman, 2018. Using stacking to average Bayesian predictive distributions. Bayesian Analysis .

[10] Vehtari, A., A. Gelman, and J. Gabry, 2017. Practical Bayesian model evaluation using leave-one-out cross-validation and WAIC. Statistics and Computing 27:1413–1432.

[11] Vehtari, A., J. Gabry, Y. Yao, and A. Gelman, 2018. loo: Efficient leave-one-out cross-validation and WAIC for Bayesian models. https://CRAN.R-project.org/package=loo, r package version 2.0.0.

[12] Gelman, A., J. Hwang, and A. Vehtari, 2014. Understanding predictive information criteria for Bayesian models. Statistics and Computing 24:997–1016.

[13] Wolpert, D. H., 1992. Stacked generalization. Neural Networks 5:241–259.

[14] Breiman, L., 1996. Stacked regressions. Machine Learning 24:49–64.

[15] Alsous, J. I., P. Villoutreix, A. M. Berezhkovskii, and S. Y. Shvartsman, 2017. Collective growth in a small cell network. Current Biology 27:2670–2676.e4.

[16] Little, S. C., M. Tikhonov, and T. Gregor, 2013. Precise developmental gene expression arises from globally stochastic transcriptional activity. Cell 154:789–800.

[17] Schindelin, J., I. Arganda-Carreras, E. Frise, V. Kaynig, M. Longair, T. Pietzsch, S. Preibisch, C. Rueden, S. Saalfeld, and B. Schmid, 2012. Fiji: an open-source platform for biological-image analysis. Nature Methods 9:676.
